# Genetic mapping of lifespan and mitochondrial stress response in *C. elegans*

**DOI:** 10.1101/2025.06.26.661693

**Authors:** Xiaoxu Li, Weisha Li, Arwen W. Gao, Yunyun Zhu, Elena Katsyuba, Katherine A. Overmyer, Terytty Yang Li, Ziwen Wang, Luc Legon, Lucie Plantade, Laurent Mouchiroud, Matteo Cornaglia, Hao Li, Riekelt H. Houtkooper, Joshua J. Coon, Johan Auwerx

**Affiliations:** Laboratory of Integrative Systems Physiology, Interfaculty Institute of Bioengineering, École Polytechnique Fédérale de Lausanne, Lausanne, Switzerland; Laboratory Genetic Metabolic Diseases, Amsterdam UMC, Location University of Amsterdam, Meibergdreef 9, Amsterdam, The Netherlands; Amsterdam Gastroenterology Endocrinology and Metabolism Institute, Amsterdam, The Netherlands; Department of Biomolecular Chemistry, University of Wisconsin, Madison, WI 53506, USA; National Center for Quantitative Biology of Complex Systems, Madison, WI 53706, USA; Morgridge Institute for Research, Madison, WI 53515, USA; Department of Chemistry, University of Wisconsin, Madison, WI 53506, USA; Nagi Bioscience SA, EPFL Innovation Park, 1025 Saint-Sulpice, Switzerland; Center for Mitochondrial Biology and Medicine, The Key Laboratory of Biomedical Information Engineering of Ministry of Education, School of Life Science and Technology, Xi’an Jiaotong University, Xi’an, China; State Key Laboratory of Genetic Engineering, Shanghai Key Laboratory of Metabolic Remodeling and Health, Laboratory of Longevity and Metabolic Adaptations, Institute of Metabolism and Integrative Biology, Fudan University, Shanghai, China

**Keywords:** *C. elegans*, UPR^mt^, systems genetics, multi-omics integration, and aging

## Abstract

The mitochondrial unfolded protein response (UPR^mt^) is one of the mito-nuclear regulatory circuits that restores mitochondrial function upon stress conditions, promoting metabolic health and longevity. However, the complex gene interactions that govern this pathway and its role in aging and healthspan remain to be fully elucidated. Here, we activated the UPR^mt^ using doxycycline (Dox) in a genetically diverse *C. elegans* population comprising 85 strains and observed large variation in Dox-induced lifespan extension across these strains. Through multi-omic data integration, we identified an aging-related molecular signature that was partially reversed by Dox. To identify the mechanisms underlying Dox-induced lifespan extension, we applied quantitative trait locus (QTL) mapping analyses and found one UPR^mt^ modulator, *fipp-1*/*FIP1L1*, which was functionally validated in *C. elegans* and humans. In the human UK Biobank, *FIP1L1* was associated with metabolic homeostasis, underscoring its translational relevance. Overall, our findings demonstrate a novel UPR^mt^ modulator across species and provide insights into potential translational research.

## Introduction

Mitochondria, essential organelles for cellular energy homeostasis, are important determinants of whole-body metabolism, health, and lifespan^1,2^. The mitochondrial matrix is an inherently stressful environment for proteins for many reasons^3^. Not only are mitochondria the seat of most metabolic transformations and susceptible to oxidative stress but they are also exposed to proteotoxic stress as the majority of mitochondrial proteins are translated in the cytosol and imported in the unfolded form to the matrix, where they are refolded and assembled^4^. Mitochondria hence evolved numerous regulatory circuits to maintain homeostasis and enable adaptation to mitochondrial stress through processes ranging from biogenesis, dynamics, mitophagy, and mitochondrial protein quality control^5^. These mitochondrial stress response (MSR) pathways rely on mito-nuclear communication, combining anterograde and retrograde signals in interlocked feedback circuits, typified by the mitochondrial unfolded protein response (UPR^mt^)^5–7^. These stress circuits are evolutionarily conserved^8–11^ and linked to a broad spectrum of diseases^3,12,13^ as well as longevity^14–18^. While the UPR^mt^ was discovered in mammalian cells^19–21^, the UPR^mt^ signal transduction has been mainly mapped in *C. elegans* through gain-/loss-of-function (G/LOF) studies^13^.

However, effects of mutations and environmental perturbations are also often—as was the case for the studies on the UPR^mt^—only examined on a single genetic background, which limits their translational utility^22^. One solution to this problem is to use worm genetic reference populations (GRPs)^23–29^. These panels consist of inbred strains derived from crosses between two genetically divergent parental strains^28,30^. In contrast to G/LOF studies, most of the effects of variation in GRPs are subtle and quantitative in nature^24^. Furthermore, the availability of the genotype data, and the ability to reproduce identical individuals allow for the in-depth interrogation of quantitative traits at the systems level in several environmental conditions and at multiple physiological levels. We have previously reported the presence of strain-specific regulations of the MSR in wild-type N2 and CB4856 worms^31^ by integration of multi-omics data analysis. Given the important role of MSR in disease pathogenesis^32^ and lifespan extension^15^, continued research into uncovering the mechanisms regulating MSR remains essential.

Here, we used 85 recombinant intercross advanced inbred lines (RIAILs) derived from crosses between two parental strains, i.e. QX1430 (with an N2 Bristol background) and CB4856 (Hawaii)^28^. To identify the alleles underlying the subtle variations in phenotypes linked with the UPR^mt^ activation across the worm GRP (**Fig. 1**), we collected samples of these strains after induction of the UPR^mt^ with doxycycline (Dox), an antibiotic that not only interferes with bacterial but also with mitochondrial translation^15^. Combining our previously analyzed control data^33^, we evaluated the effect of Dox on lifespans and life-history phenotypes—including body size, developmental dynamics, and reproductive/fertility parameters—as well as on multi-omics datasets (transcriptome, proteome, metabolome, and lipidome). In addition, the integration of these omics data and lifespan allowed the indication of longevity-related molecular features and the identification of genetic loci responsible for UPR^mt^ regulators. We identified and validated *fipp-1*, which encodes factor interacting with Poly(A) polymerase, as a regulator of the UPR^mt^ in *C. elegans*. We then translated these findings to human cells and demonstrated that its ortholog, *FIP1L1*, plays a critical role in regulating the MSR pathway. Moreover, by integrating data from large-scale human cohorts—including the UK Biobank (UKBB) and the Million Veterans Program (MVP)— we found that genetic variants within *FIP1L1* are associated with metabolic health homeostasis in humans.

**Figure 1:**
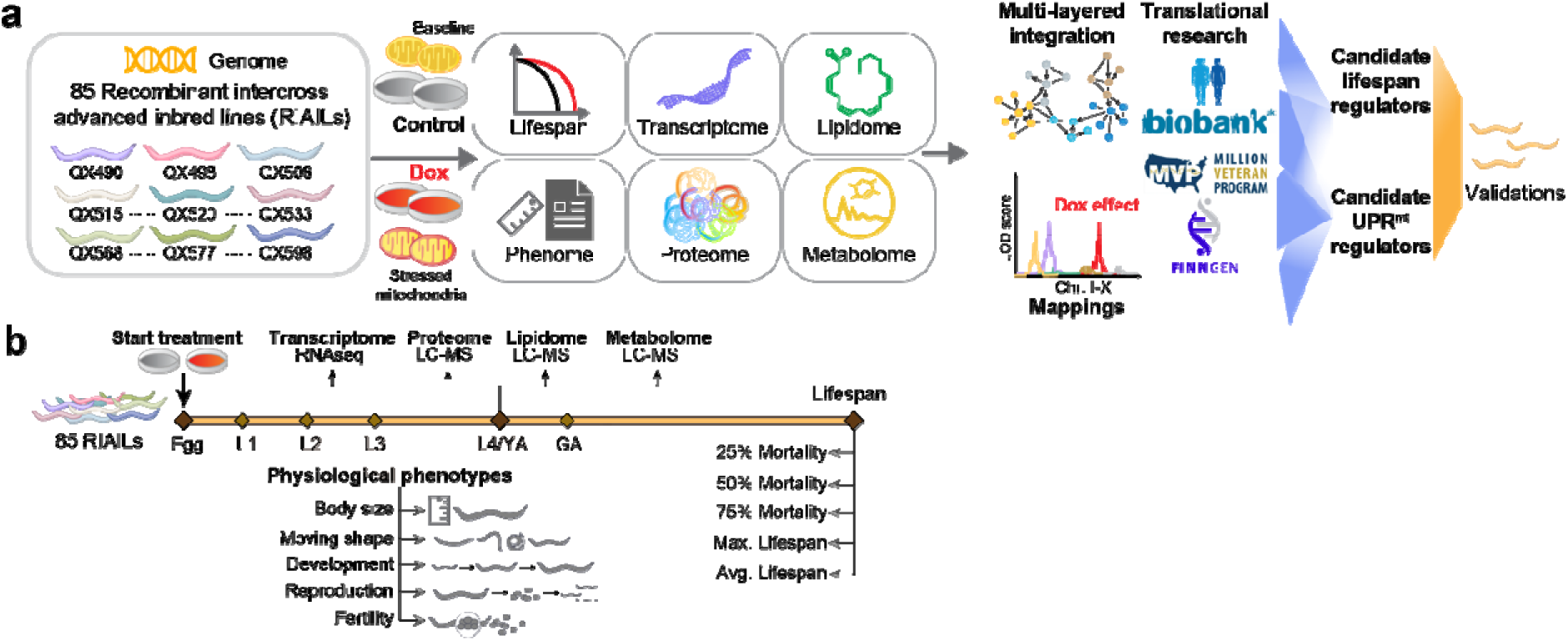
Overview of the study. (**a**) 85 recombinant intercross advanced inbred lines (RIAILs) derived from crosses of QX1430 (N2 Bristol background, with deletions of confounder genes) and CB4856 strain (Hawaii) were used to measure lifespan, early life-history traits, transcriptome, lipidome, proteome, and metabolome. We applied system genetics approaches to identify candidate genes for regulating lifespan and mitochondrial stress response (MSR) induced by Dox. After prioritization of the candidate genes, they were first explored in the human large-scale databases and then validated with wet lab experiments. (**b**) Experimental procedures and sampling. All the data are collected from three groups of RIAILs. One group of RIAILs was cultured and scored for lifespan; one group of RIAILs was cultured in th microfluidic device for ∼100 h to collect early life-history traits, including body size, moving shapes, developmental parameters, reproduction, and fertility; and the last group of RIAILs was cultured to reach L4/young adulthood, and collected for RNA-sequencing (RNA-seq), proteomics, lipidomics, and metabolomics measurements. Chr. I-X: chromosome I to X; LC-MS: liquid chromatography – mas spectrometry; L1-L4: larval stage 1 to 4; YA: young adulthood; GA: gravid adulthood. Max. Lifespan: maximum lifespan; Avg. Lifespan: average lifespan.

## Results

### Dox affects the lifespan and phenotypic traits of 85 RIAILs in a strain-specific manner

To assess the effect of Dox-induced MSR on lifespans in 85 RIAILs, we first performed survival analysis and extracted five commonly used traits^33^ from the survival curves, including 25% mortality (defined as the day when 25% of worms of each strain were dead), 50% mortality, 75% mortality, maximum (max.) lifespan and average (avg.) lifespan. In a global and strain-independent manner, we confirmed the Dox effect on lifespan extension and observed an overall beneficial effect of Dox at all stages of worm life (**Fig. 2a-b**). We also estimated the Dox effect in a strain-dependent view and found large variations in their lifespan extension upon Dox exposure, represented by hazard ratio (HR) (**Fig. 2c and S1**). Most RIAILs have extended lifespans upon Dox (e.g. QX594, indicated in green), whereas some strains have either shortened (e.g. QX585, colored in red) or unchanged (e.g. QX590, represented by grey) lifespans (**Fig. 2c and S1**). Given the large variation of lifespan we previously observed in the control condition^33^, our result indicated that genetic background not only plays an essential role in shaping lifespan but also affects the response to Dox exposure.

**Figure 2:**
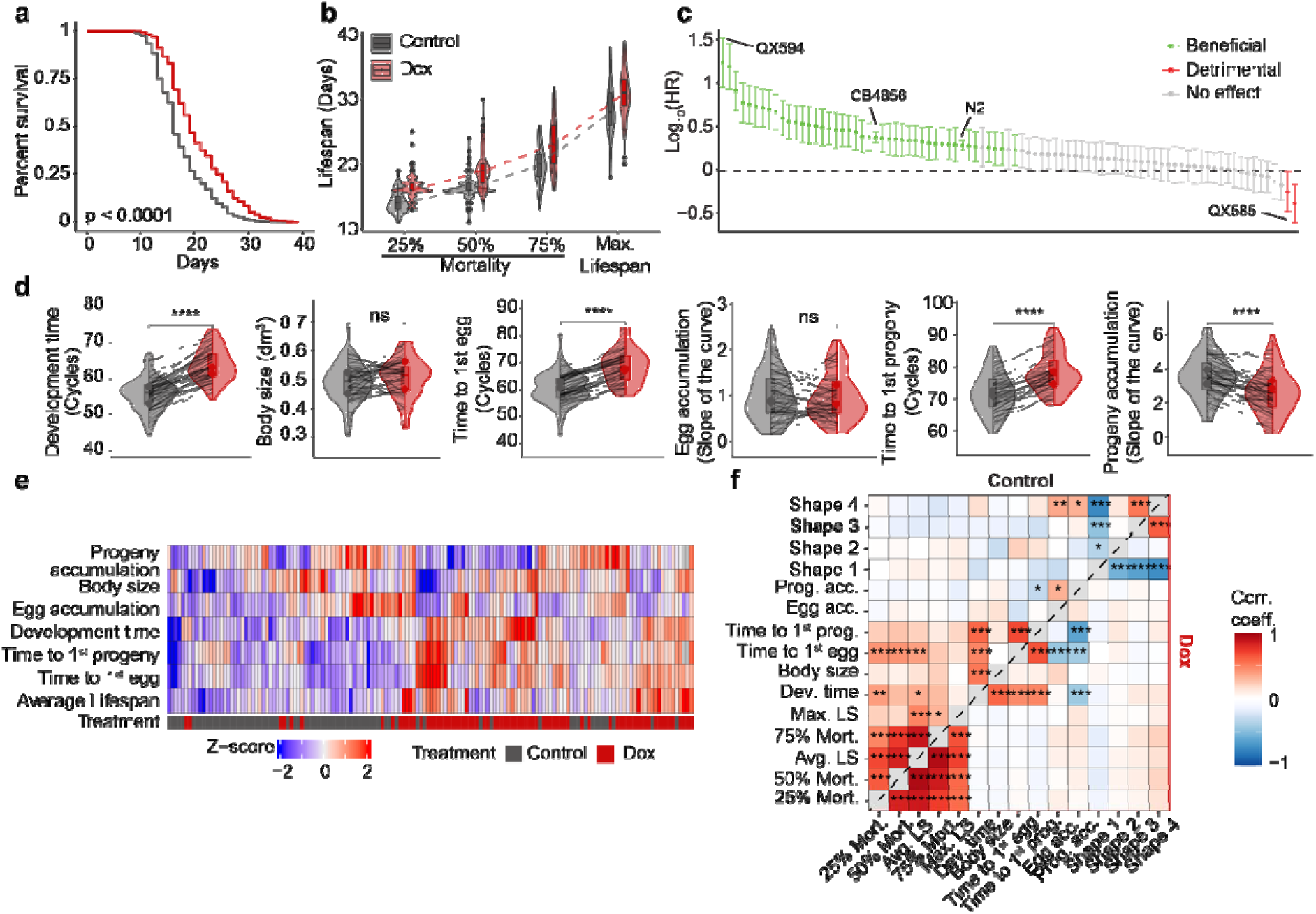
Overview of Dox effect on longevity and phenotypes. **(a)** Kaplan–Meier curve showed that Dox extended the lifespan of the RIAILs population. Control condition: black lines; Dox treated: red line. (**b**) Boxplots and distributions of 25%, 50%, and 75% survival as well as maximal lifespan under control (black) and Dox (red) conditions. **(c)** Lifespan extension by Dox was calculated by -Log_10_(Hazard ratio). Each column indicates a strain of RIAILs. Solid circles represent the effect, and the bars the associated 95% confidence interval. The dashed line indicates no effect on lifespan. Green: RIAILs with extended lifespans; Grey: non-significant; red: RIAILs with reduced lifespans upon Dox. (**d**) Violin plots of life-history phenotypic traits under control (black) and Dox (red) conditions. P-values represent the comparison of each trait between the control and Dox, calculated using a two-tailed Student’s t-test. Line indicate the changes of each phenotype in each strain upon Dox. (**e**) Hierarchical clustering of life-history phenotypic traits and average lifespan. (**f**) Pearson correlation between lifespan and physiological traits under control (upper-left) and Dox-treated (bottom-right) conditions.

In addition to the effects of environmental exposures and genetics, lifespan displays significant associations with clinical phenotypic traits in mice and humans. To further investigate how Dox exposure affects early-life phenotypic traits and their association with lifespan, we monitored these traits under Dox exposure and compared them to those previously characterized under untreated conditions^33^. Principal component analysis (PCA) revealed that the effects of Dox on phenotypic traits were primarily captured by PC1 **(Fig. S2a)**. At the population level, the developmental time was extended by the mitochondrial stress induced by Dox, suggesting that the whole population has an overall developmental delay when exposed to Dox (**Fig. 2d**). However, the body size that the worms reached (**Fig. 2d**) and most activities that the worms performed in liquid (**Fig. S2b-c**) were not affected by Dox exposure. In line with a slower growth dynamic, Dox delayed the sexual maturity of the RIAILs (capacity to lay eggs) and their progeny appearance as well as reduced progeny accumulation (**Fig. 2d**). These data confirmed that the retarded development is an evolutionary cost of lifespan extension upon Dox, which is consistent with our previous findings^33^ that slow growth is positively correlated with longevity in *C. elegans*.

However, similar to the changes in lifespan, we also found a variable Dox effect on clinical phenome in a strain-specific manner (**Fig. 2e and S2b-d**). To illustrate whether collected phenotypic traits are associated with longevity after Dox exposure, we applied Pearson correlation analysis and found that the positive association between longevity and slow development are eliminated under Dox exposure **(Fig. 2f)**. This suggests that the effect of Dox outweighs the influence of the detected phenotypes on longevity.

### Exploration of the effects of Dox on multi-omics datasets

To investigate the molecular phenotypic changes induced by Dox, we assessed multi-omics datasets, including transcriptomics, proteomics, metabolomics, and lipidomics of RIAILs exposed to Dox treatment in the population level. By integrating our previously analyzed baseline condition datasets^33^, we conducted differential expression analysis and observed that the changes in gene and protein expression upon Dox are partially consistent (**Fig. 3a** and **S3a-b**, correlation coefficient = 0.59). The significantly upregulated and downregulated overlapping genes and proteins revealed global regulatory responses to mitochondrial stress. Consistent with findings in wild-type *C. elegans* strains^31^, genes involved in innate immune response and the endoplasmic reticulum (ER) UPR were upregulated at both transcriptomic and proteomic levels following Dox exposure, while genes associated with DNA replication and cell cycle progression were downregulated (**Fig. 3b)**. Additionally, we assessed strain-specific UPR^mt^ responses to Dox and confirmed that Dox consistently elevated UPR^mt^-related protein levels in RIAIL strains (**Fig. 3c**). We also observed a variation in Dox-induced effects among these RIAILs (**Fig. 3c**), which may be attributed to genetic differences.

**Figure 3:**
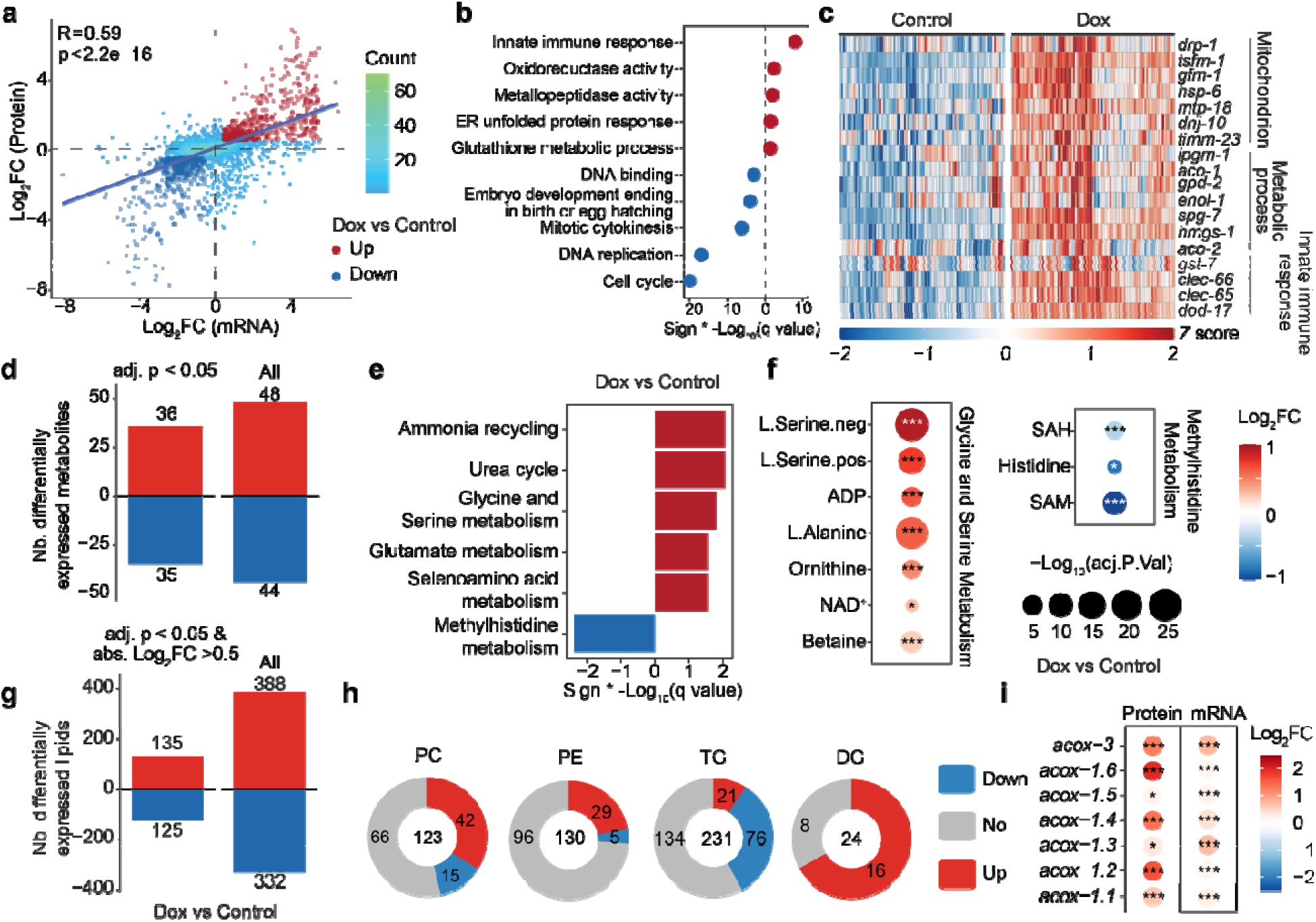
The effect of Dox on multi-omic traits in the RIAILs. (**a**) Scatter plot showing the Dox effect on mRNA-protein pairs at the mRNA level (x-axis) and protein level (y-axis). Overlapped up- and down-regulated genes (adjusted p-value < 0.05, Log_2_(fold-change) > 0.5 or < −0.5) induced by Dox are indicated by red and blue, respectively. Differential gene expression upon Dox treatment, irrespective of strains. **(b)** Dot plot showing the enriched gene sets of overlapped up-(indicated in red) and down-regulated gene (indicated in blue) upon Dox. **(c)** Heatmap displays the changes of UPR^mt^-related proteins upon Dox. **(d)** Bar plot indicating the impact of Dox on the metabolome. **(e)** Metabolite pathway enrichment analysis of up-(36 metabolites, indicated in red) and down-regulated metabolites (35 metabolites, indicated in blue) upon Dox. **(f)** The changes of metabolites that are involved in glycine and serine metabolism, as well as methylhistidine metabolism. Log_2_FC is represented by color and the -Log_10_(adj.P.Val) is shown by dot size and indicated as * -Log_10_(adj.P.Val) < 0.05, ** -Log_10_(adj.P.Val) < 0.01, *** -Log_10_(adj.P.Val) < 0.001. **(g)** Barplot indicating the impact of Dox on the lipidome. **(h)** Pie plots showing the effect of Dox on PC, PE, TG, and DG lipid classes. **(i)** Dox effects on the genes or proteins involved in the fatty acid beta oxidation. Log_2_FC is represented by color and the -Log_10_(adj.P.Val) is shown by dot size and indicated as * -Log_10_(adj.P.Val) < 0.05, ** -Log_10_(adj.P.Val) < 0.01, *** -Log_10_(adj.P.Val) < 0.001.

In addition, the impact of the Dox-induced MSR on metabolomics and lipidomics of the RIAILs was assessed in a strain-independent manner. Among the measured metabolites, 36 were significantly elevated, and 35 were significantly reduced (**Fig. 3d**). Functional clustering of the altered metabolites revealed enrichment in several key metabolic pathways. Specifically, metabolites associated with ammonia recycling, the urea cycle, glycine and serine metabolism, glutamate metabolism, and selenoamino acid metabolism were significantly increased upon Dox exposure (**Fig. 3e**). In contrast, those involved in methylhistidine metabolism were decreased (**Fig. 3e**). Of note, we observed significant elevations in serine, alanine, ornithine, ADP, NAD^+^, and betaine levels (**Fig. 3f**). Increased serine suggests activation of one-carbon metabolism, crucial for nucleotide synthesis and methylation processes associated with lifespan extension^34–36^. Elevated alanine levels indicate alterations in amino acid metabolism typically seen as metabolic signatures of long life in *ife-2(ok36)* and *daf-2(m41)* mutants^37^. Elevated ADP levels are consistent with disrupted energy homeostasis resulting from mitochondrial dysfunction induced by Dox. The observed increase in NAD^+^ aligns with previous findings of enhanced mitochondrial function and lifespan extension^38^, suggesting that Dox-induced UPR^mt^ reshapes NAD^+^ metabolism and it in turn contributes to the observed stress resistance and longevity associated with the UPR^mt^. Additionally, elevated betaine levels reflect enhanced longevity mechanisms, potentially through the promotion of autophagy and antioxidant activities^39^. Conversely, several metabolites were significantly reduced following mitochondrial stress induction by Dox, including S-adenosylmethionine (SAM), S-adenosylhomocysteine (SAH), and histidine (**Fig. 3e-f**). Reduced SAM and SAH levels suggested disrupted methylation homeostasis, a phenomenon associated with delayed aging and lifespan extension in *C. elegans*^40^. Together, these metabolic changes represent adaptations consistent with mitochondrial stress-induced metabolic reprogramming, highlighting pathways linked to MSR and longevity regulation. Furthermore, Dox treatment significantly altered lipid levels, with 135 increasing and 125 decreasing (**Fig. 3g**). Lipidomic profiling by lipid class revealed that Dox significantly decreased most triglycerides (TGs) levels, whereas phosphatidylcholine (PC), phosphatidylethanolamine (PE), and diglycerides (DGs) levels were increased in response to mitochondrial stress (**Fig. 3h**). Consistent with this observation, key genes involved in beta-oxidation, including acyl-CoA oxidases (ACOXs), were upregulated following Dox exposure (**Fig. 3i**) —likely as a compensatory response to increased fatty acid availability from Dox-induced TG hydrolysis. These findings suggest that Dox promotes lipolysis, elevating substrate flux into beta-oxidation and driving a metabolic shift in lipid homeostasis under mitochondrial stress.

### Identification of lifespan regulators from the transcriptome of RIAILs

Intermediate molecular phenotypes play a crucial role in modulating aging^41^. To identify lifespan-associated genes within the transcriptome and elucidate key gene regulators influencing longevity, we first constructed co-expressed gene modules using weighted gene co-expression network analysis (WGCNA) based on transcriptome data collected in the control and Dox condition. Then, we applied Cox regression analysis to estimate lifespan-related gene modules **(Fig. 4a)**. Our analysis revealed that higher eigengene values in Modules 5, 13, and 14 were significantly associated with increased lifespan (BH-adjusted p-value < 0.05, -Log□□(HR) > 0, **Fig. 4b**), suggesting that these modules serve as potential predictors of longevity.

**Figure 4:**
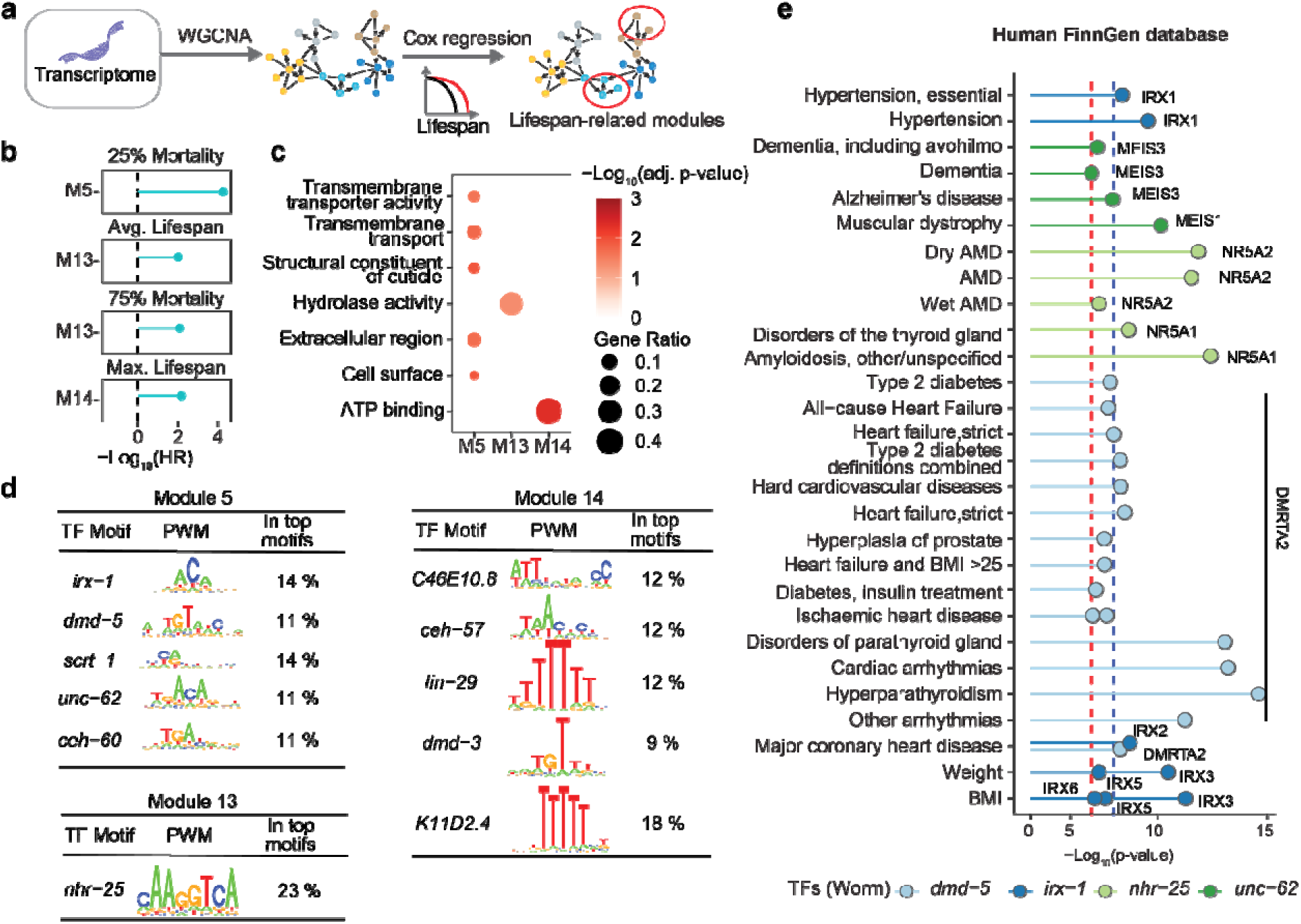
Biological interrogation of lifespan trait-related gene modules constructed from the transcriptome. **(a)** Schematic of the pipeline used to identify lifespan trait-associated gene modules from transcriptomic data. Gene co-expression networks were constructed by WGCNA, and Cox regression was applied to evaluate lifespan-related modules. **(b)** -Log_10_(Hazard ratio) of mixed-effects Cox regression model for lifespan traits, with the eigengenes of gene modules as predictor values, Dox and the interaction between phenotype and Dox as covariates, and RIAIL strain as random effect. Only gene modules that were significantly associated with lifespan traits after multiple testing adjustments with the BH method (BH-adjusted p value < 0.05) are shown. **(c)** Dot plot showing the enriched pathways for genes involved in lifespan-related modules. **(d)** Enriched transcription factor binding motifs in the promoters of genes within Modules 5, 13, and 14. **(e)** Dot plot showing associations between enriched transcription factors and metabolic disorders in the human FinnGen database. Only suggestive (-Log_10_P > 7, red dashed line) or significant (-Log_10_P > 8, blue dashed line) associations are shown. AMD: Age related macular degeneration.

To further explore the biological relevance of these modules, we performed a functional clustering analysis. Genes within Module 13 were predominantly linked to hydrolase activity, whereas those in Module 14 were associated with ATP binding and Module 5 genes were enriched for transmembrane transporter activity **(Fig. 4c)**. Additionally, we identified transcription factors (TFs) that regulate gene expression within each module. The top five TFs modulating Module 5 were *irx-1*, *dmd-5*, *scrt-1*, *unc-62*, and *ceh-60*. For Module 14, key regulators included *C46E10.8*, *ceh-57*, *lin-29*, *dmd-3*, and *K11D2.4*, while *nhr-25* was the only TF significantly associated with Module 13 **(Fig. 4d)**. These findings provide insights into the regulatory mechanisms underlying transcriptomic influences on longevity.

Next, to assess whether the identified TFs also play a significant role in human health, we examined seven TFs with human orthologs using human genetic analyses. By integrating data from the FinnGen database^42^, we explored associations between genetic variants within these TFs and various phenotypic traits. Our analysis revealed that genetic variants in the human orthologs of *dmd-5* (*DMRTA2* and *DMRTB1*) and *irx-1* (Iroquois homeobox (IRX) protein family genes (*IRX1-6*)) were associated with metabolic traits, including body weight, cardiovascular disease, and type 2 diabetes (**Fig. 4e**). Additionally, the orthologs of *nhr-25* (*NR5A1* and *NR5A2*) and *unc-62 (MEIS1, MEIS2, MEIS3)* were linked to aging-related diseases such as dementia and Alzheimer’s disease (**Fig. 4e**). These findings further underscore the importance of these TFs in modulating human health.

To experimentally validate the role of these TFs in lifespan regulation, we performed RNAi-mediated knockdown of *dmd-5*, *irx-1*, and *nhr-25* in worms and assessed their effects on longevity. Intriguingly, worms subjected to *dmd-5* or *irx-1* RNAi exhibited significantly extended lifespans (**Fig. S4a-b**), whereas *nhr-25* RNAi resulted in lifespan shortening **(Fig. S4c**). Furthermore, *unc-62*, a known aging-related gene, has been confirmed to promote longevity upon knockdown in a previous study^43^. Collectively, by integrating *C. elegans* transcriptomic data with genome-wide association study (GWAS) findings from a large-scale human dataset, we identified four TFs that regulate lifespan in *C. elegans* and influence human health.

### Exploring the molecular signature of longevity by integrating multi-omics datasets

While individual omics datasets provide valuable insights into lifespan and metabolic health, multi-omics profiling enables the integration of distinct biological signals across complementary layers, facilitating a more comprehensive understanding of molecular interconnections and the identification of system-level biomarkers. In this study, we applied multi-omics factor analysis (MOFA)^44^ to integrate proteomic, metabolic, and lipidomic datasets, reducing their dimensionality into ten factors under both control and Dox-treated conditions (**Fig. 5a**). The eigengene values of these ten factors were subsequently used as predictors in a Cox regression model to evaluate their prognostic value for five lifespan-related traits, with Dox treatment included as a confounder (**Fig. 5a**).

**Figure 5:**
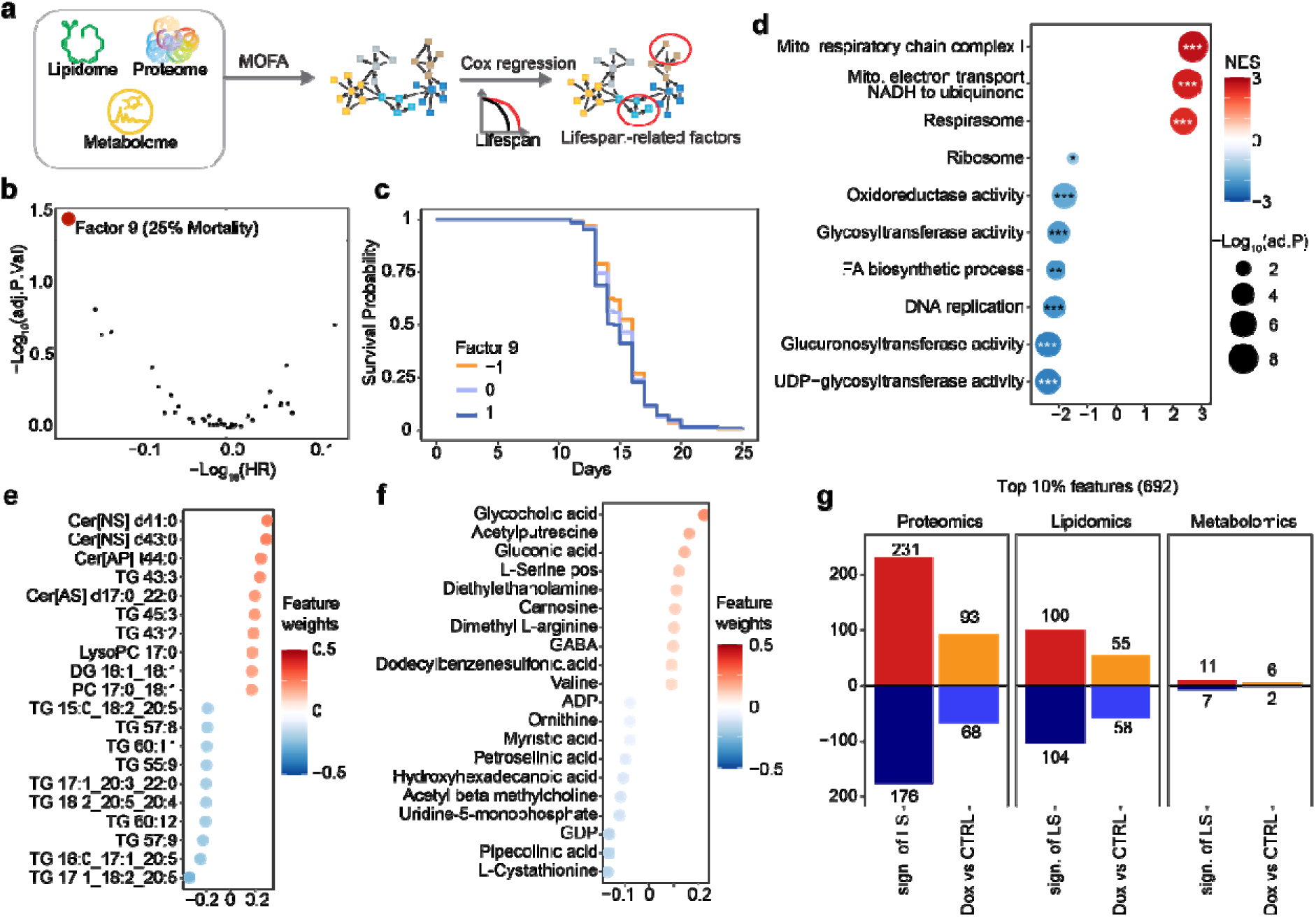
Identification of a longevity signature in the proteome, metabolome, and lipidome. (**a**) Schematic of the pipeline used to explore lifespan-associated factors across multi-omics data, including proteome, metabolome, and lipidome. (**b**) Volcano plot displaying the association between factor value and lifespan traits evaluated by the Cox regression model. Red dots indicate factors significantly associated with lifespan (adjusted P value < 0.05). (**c**) Kaplan–Meier curve shows that the eigengene value of factor 9 is negatively associated with 25% mortality. Orange, light blue, and dark blue color indicate the eigengene value < −1, between −1 to 1, and > 1, respectively. (**d-f**) Dot plot showing top features contributing to factor 9 in proteomic (**d**), lipidomic (**e**), and metabolomic (**f**) analysis. Given that a higher eigengene value of factor 9 correlated with reduced lifespan, we inverted these weights by multiplying them by −1 to obtain feature weights that were positively associated with longevity. (**g**) Bar plots display the number of top 10% features that positively or negatively influence lifespan extension, along with how many of these features are affected by Dox exposure. Sign. of LS: the lifespan-associated signatures -- the top 10% of features contributing to longevity in factor 9 (629/6292 features). Dox v CTRL: the Dox-induced changes in proteins, lipids, and metabolites.

Notably, we identified factor 9 as negatively associated with the lifespan of 25% mortality (BH-adjusted p-value < 0.05, -Log□□(HR) < 0; **Fig. 5b-c**). To further investigate the molecular features linked to lifespan, we extracted the feature weights of proteins, metabolites, and lipids contributing to factor 9. Given that a higher eigengene value of factor 9 correlated with reduced lifespan, we inverted these weights by multiplying them by −1 to obtain feature weights that were positively associated with longevity. Using the weights of proteins in factor 9, gene set enrichment analysis (GSEA) suggested that an increased abundance of proteins involved in mitochondrial function was positively associated with longevity, whereas proteins related to fatty acid biosynthesis and cell proliferation were negatively correlated with lifespan (**Fig. 5d**). Consistent with these proteomic findings, our lipidomic analysis revealed that most triglycerides (TGs) negatively contributed to lifespan, including TG 17:1_18:2_20:5, whereas two non-hydroxy-fatty acid [N] sphingosine [S] ceramides were positively associated with longevity (**Fig. 5e**). Additionally, glycocholic acid (GCA) and N-acetylcysteine (NAC) emerged as the top two metabolites positively linked to lifespan, both of which have been previously reported to exhibit anti-inflammatory and antioxidant properties^45,46^ (**Fig. 5f**). Collectively, these findings suggest that longevity is associated with enhanced mitochondrial function, elevated levels of GCA and NAC, and reduced TG abundance.

Given that Dox treatment extends lifespan, we next investigated whether its effects on molecular phenotypes aligned with the lifespan-associated signatures we identified. We first extracted the top 10% of features contributing to longevity in factor 9 (629/6292 features) as the lifespan-associated signatures (Sign. of LS). The Dox-induced changes in proteins, lipids, and metabolites were then identified. By comparing Dox-induced changes in these omics’ datasets to the identified Sign. of LS, we found that nearly half of the lifespan-associated features exhibited consistent changes upon Dox exposure (**Fig. 5g**). These results provide mechanistic insights into how Dox influences molecular pathways to promote longevity.

### Uncovering genetic determinants regulating MSR induced by Dox

In addition to the exploration of Dox effects on molecular traits, we next sought to uncover how genetic background affects the lifespan extension induced by MSR. We performed quantitative trait locus (QTL) mapping across 85 wild-derived strains and identified a significant QTL peak on chromosome V, indicating that genetic variation at this locus modulates the response to Dox (**Fig. 6a**). Notably, strains inheriting the CB4856 allele at this locus exhibited significantly greater lifespan extension upon Dox treatment compared to those with the N2 allele (**Fig. 6b**). To further explore candidates under this QTL peaks, we applied three criteria **(Fig. 6c)**: (1) having known segregating variants in the parental strains predicted to have high/moderate impact, (2) having a *cis*-expression QTL (*cis*-eQTLs), indicating that the expression of the gene is controlled by the same locus, and (3) differentially expressed genes (DEGs) upon Dox in transcriptome. Collectively, 13 genes satisfying two of the above criteria were considered candidate genes that might influence the Dox effect on lifespan extension in *C. elegans* **(Fig. 6d)**.

**Figure 6:**
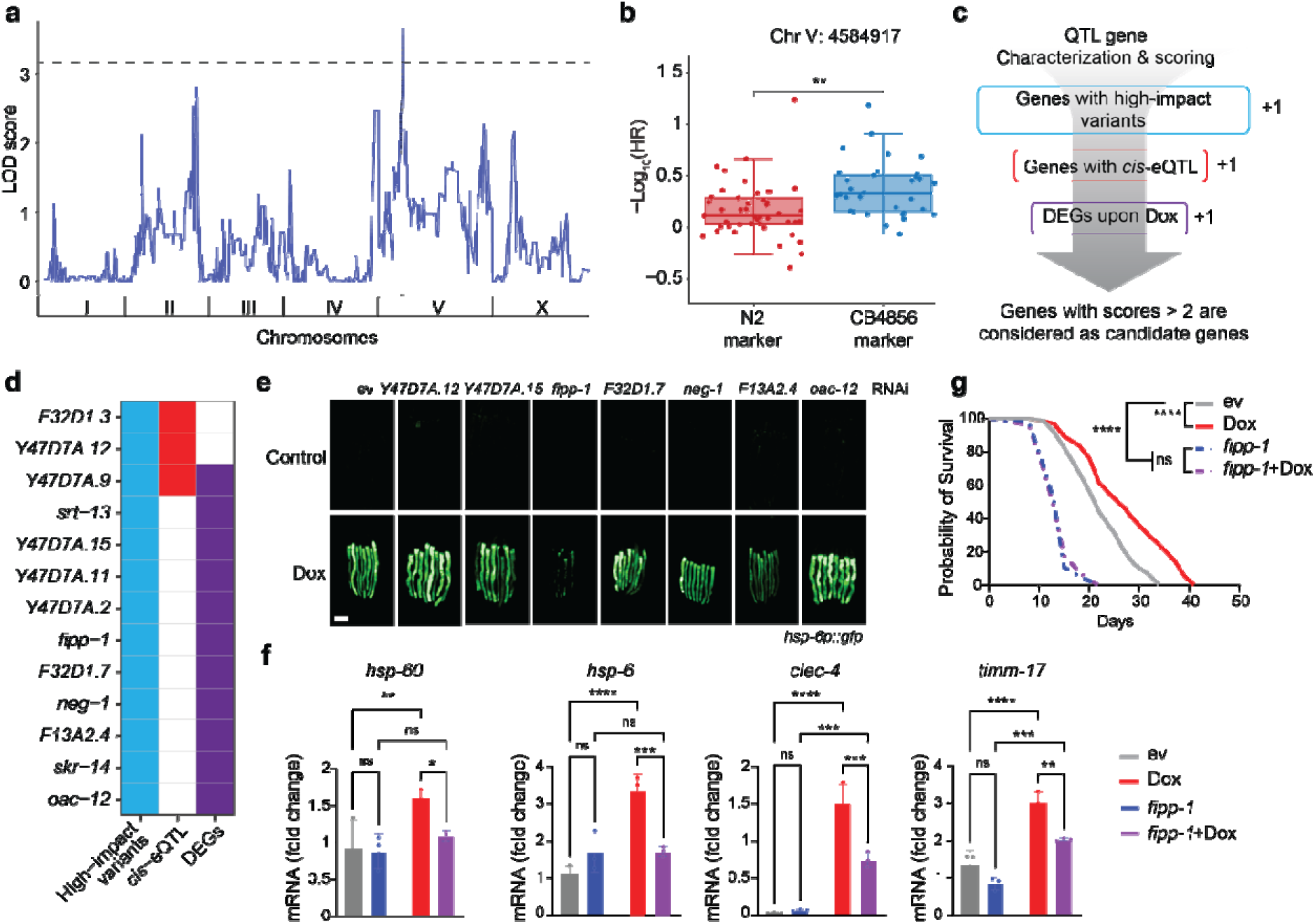
Prioritizing MSR regulators in *C. elegans.* (**a**) Manhattan plot showing the QTL mapping result of Dox-induced lifespan extension. (**b**) Boxplot showing the effect of the genetic locus located on Chr. V on Dox-induced lifespan extension. (**c**) A schematic indicating the identification and prioritization of candidate genes regulating MSR-induced lifespan extension. DEGs: differentially expressed genes. (**d**) The identified candidates under this QTL peak are according to the selection criteria in (c). (**e**) GFP expression levels of *hsp-6p::gfp* worms treated with RNAi targeting different candidate genes (*Y47D7A.12*, *Y47D7A.15*, *fipp-1*, *F32D1.7*, *neg-1*, *F12A2.4*, and *oac-12*) with or without Dox treatment. scale bar: 150µm. (**f**) qRT-PCR analysis of transcripts (n= 3 biological replicates) in worms treated with control (ev), or RNAi targeting *fipp-1*, treated with or without Dox. (**g**) Survival curve of worms treated with *fipp-1* RNAi with or without Dox. Statistical analysis was performed by the log-rank test. ns: not significant; ****P<0.0001; **P<0.01; *P<0.05.

To investigate the role of candidate genes in MSR regulation, we knocked down each gene individually by feeding worms with RNAi-expressing bacteria, beginning at the maternal stage. However, for six of these candidate genes, RNAi clones were not available in existing libraries. We did not generate new constructs, as transcriptomic data revealed their extremely low expression levels in *C. elegans* **(Fig. S3c)**, making RNAi-mediated knockdown technically impractical. Therefore, we proceeded with only the seven genes that had correct RNAi constructs for further investigation. We used the *hsp-6p::gfp* reporter strain to assess whether these candidate genes influence the activation of the UPR^mt^ upon exposure to Dox. We observed that RNAi-mediated knockdown of *fipp-1* (homologous to human *FIP1L1*, factor interacting with *PAPOLA* and *CPSF1*) strongly suppressed GFP induction in *hsp-6p::gfp* worms following Dox exposure (**Fig. 6e**). We further analyzed the expression of additional UPR^mt^-associated genes and observed that knockdown of *fipp-1* not only blocked the induction of mitochondria-related genes (*hsp-6* and *hsp-60*), but also significantly abrogated the induction of innate immune response-related transcripts, such as *clec-4* (**Fig. 6f**), suggesting that *fipp-1* plays a critical role in coordinating mitochondrial stress signaling with innate immunity. Importantly, RNAi of *fipp-1* completely abolished the lifespan extension induced by Dox (**Fig. 6g**), further suggesting that the activation of innate immune pathways mediated by *fipp-1* may be essential for the beneficial longevity effects observed during mitochondrial stress.

### Exploring the biological function and clinical relevance in mice and humans

To understand the potential conserved roles of this validated UPR^mt^ regulator, we studied its homolog, *FIP1L1*, in humans. Dox was used to induce the UPR^mt^ in human embryonic kidney (HEK) 293T cells with or without *FIP1L1* knockdown. At the protein level, Dox mildly increased EIF2α phosphorylation, ATF4 and ASNS expression^47^ **(Fig. 7a)**. Interestingly, these inductions were abolished upon *FIP1L1* knockdown **(Fig. 7a)**, consistent with our findings in *C. elegans* and further support the important role of *FIP1L1* in regulating MSR induced by Dox.

**Figure 7.**
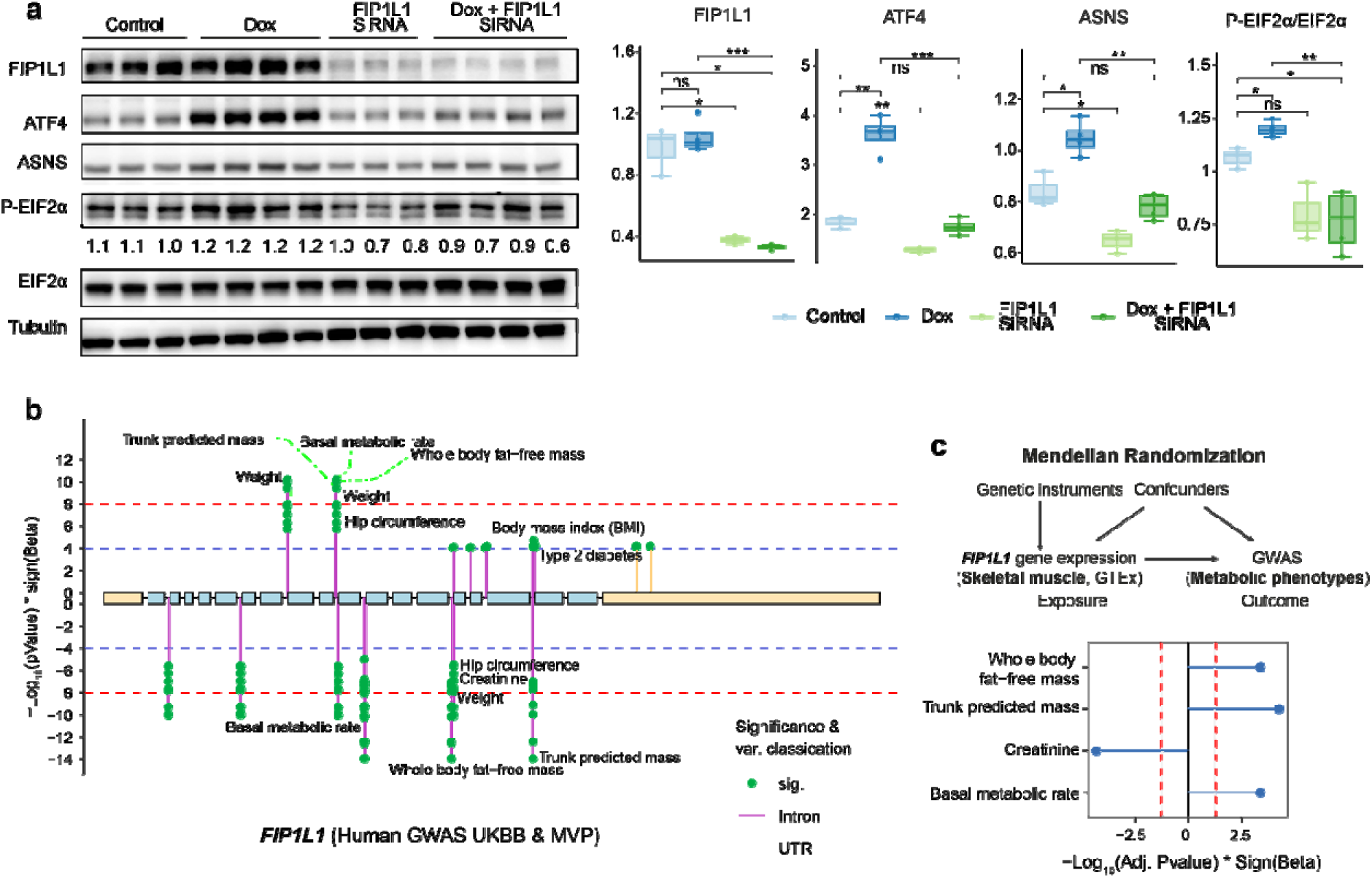
Exploring the biological function and clinical relevance across species. **(a)** Western blot analysis of UPR^mt^-related protein expression in HEK293T cells transfected with control (Ctrl) or FIP1L1 siRNA, and treated with or without doxycycline (Dox; 30Cμg/mL) for 24 hours. **(b)** Lollipop plot showing the metabolic health-related clinical traits that have suggestive (-Log10(P value) > 4) or significant ((-Log_10_(P value) > 8) GWAS hits within/around the *FIP1L1* in the UKBB and million veteran program (MVP). **(c)** Mendelian randomization analysis for the causal effect of the gene expression of *FIP1L1* in human skeletal muscle on metabolic-related traits. Thresholds were represented by red lines (BH adjusted P value < 0.05).

To investigate the eventual clinical relevance of this candidate gene, we examined phenotypic associations of genetic variants within *FIP1L1* using data from the UK Biobank (UKBB) and the Million Veteran Program (MVP). These genetic variants are significantly associated with predicted trunk mass, basal metabolic rate, as well as whole body fat-free mass (**Fig. 7b**). To assess the causal impact of *FIP1L1* expression on these traits, two-sample Mendelian randomization (MR) was performed. *cis*-eQTLs affecting *FIP1L1* gene expression were only found in one metabolic organ, i.e., skeletal muscle, in the GTEx database^48^, preventing further investigation in other metabolic tissues. In human muscles, higher gene expression of *FIP1L1* is significantly related to improved metabolic health traits, such as increased trunk predicted mass, whole body fat-free mass, and increased basal metabolic rate, and reduced creatinine levels (**Fig. 7c**). These analyses in humans further indicated the importance of *FIP1L1* in maintaining metabolic health in humans and may help explain the shortened lifespan observed in *C. elegans* following *fipp-1* knockdown.

## Discussion

Having made significant advances in dissecting individual molecules, cells, genes, and pathways contributing to aging, aging biology research is now evolving toward integrating these components to better understand how their collective dynamics contribute to longevity^49^. This shift necessitates adopting a formal complex systems framework, facilitated by large-scale datasets and advanced analytical tools. In line with this, we have identified shared and strain-specific UPR^mt^ regulators affecting lifespan extension based on multi-omics datasets in the two parental strains of the RIAILs^31^. Longevity regulators within the RIAILs under the control condition were also highlighted in our previous study using systems genetic approaches^33^. Expanding on these foundational studies, we now generated extensive physiological phenotypes and multi-omics profiles from 85 RIAIL strains cultured with doxycycline (Dox). This resulted in a comprehensive multi-omics atlas of 85 RIAILs, serving as a robust resource for investigating the regulation of mitochondrial stress and longevity.

Diverse lifespan responses among RIAILs exposed to the mitochondrial stressor, Dox, were observed. While most strains displayed increased longevity consistent with the results in the parental strains, several strains showed no lifespan change, and notably, two strains exhibited significantly reduced lifespans upon Dox exposure. Such variations underscore the crucial effect of the gene-by-Dox interaction on lifespan extension^31,50^. Dox exposure also abolished the previously observed correlation between lifespan and phenotypic traits, such as developmental time^33^. This decoupling likely reflects the dominant effects of Dox-induced mitochondrial stress overriding genetic differences that typically link phenotypic traits to longevity.

The multi-layered dataset presented here enables the identification of regulatory effects at various biological scales, further enabling the understanding of the mechanism underlying Dox exposure. Our analyses confirmed that genes involved in the UPR^mt^ were broadly upregulated at transcriptional and proteomic levels following Dox treatment. We also found a large variation in the UPR^mt^ activation across strains, while this variation did not correlate strongly with that of lifespan extension. This might be because the genetic backgrounds underlie molecular susceptibilities for Dox exposure^31,50^, including the regulatory network by UPR^mt^ activation. These networks, in turn, may differentially engage co-regulated processes, such as innate immune responses^51–53^, metabolic adaptations^54^, or lipid homeostasis^31,55^, further determining whether mitochondrial stress translates into beneficial longevity outcomes. In addition, our metabolomics analysis indicated significant increases in metabolites related to glycine and serine pathways upon Dox, pointing toward adaptive metabolic shifts likely required to maintain cellular homeostasis and mitochondrial function during stress conditions^31^. Lipidomics data revealed reduced TG levels alongside increased levels of DGs, PC, and PE upon Dox. These alterations likely reflect an adaptive restructuring of lipid metabolism under mitochondrial stress, possibly promoting membrane stability or signaling functions critical to stress adaptation^56^. Moreover, NAD^+^ levels significantly increased in Dox-treated RIAILs, suggesting enhanced mitochondrial biogenesis, redox balance, or improved energetic states as potential mediators of lifespan extension under mitochondrial stress^38^.

By integrating multi-omics profiles and lifespan data under control and Dox conditions, we identified lifespan regulators and longevity signatures. In the FinnGen database, we found that four of the identified lifespan-related transcription factors—*dmd-5*, *irx-1*, *nhr-25*, and *unc-62*—are also associated with human metabolic diseases, emphasizing their potential as targets for interventions addressing age-related pathologies. Consist with the GWAS result, two of the human homologs of *irx-1*, *IRX-3* and *IRX-5*, have been reported to link with body weight. Particularly, elevated expression of IRX3/5 in mesenchymal adipocyte precursors promotes white adipocyte formation, while in the hypothalamus, it decreases energy expenditure and stimulates food intake, reducing obesity^57,58^. Furthermore, the human homologs of *nhr-25*, *NR5A1* and *NR5A2*, have also been implicated in liver damage and related disorders^59^. These evidences further highlight the translational ability of *C. elegans* multi-omics analysis. Additionally, the overlap between Dox-induced multi-layered changes and longevity signatures underscores the specific role of MSR in lifespan extension, aligning with conserved pro-longevity mechanisms.

We also identified a QTL peak associated with lifespan extension induced by MSR. Through *in vivo* experiments in *C. elegans*, *fipp-1* was found as a potential regulator modulating MSR and affecting lifespan. Consistently, the homolog of *fipp-1*, *FIP1L1*, has also shown to suppress doxorubicin (Dox)-induced MSR in human cells. Additionally, the genetic variants within *FIP1L1* were also linked to the obesity-related traits and the MR analysis revealed that its muscle gene expression is positively associated with improved health outcomes, underscoring the important roles of *FIP1L1* in human health. Furthermore, *FIP1L1* (factor Interacting with *PAPOLA* and *CPSF1*) is involved in mRNA polyadenylation **(Fig. S5)**, a critical process for mRNA stability, nuclear export, and translation^60^. Of note, the *FIP1L1-PDGFRA* fusion has been identified as a causal factor of idiopathic hypereosinophilic syndrome and chronic eosinophilic leukemia^61^. Despite that there are no direct evidence currently links *FIP1L1* to MSR, our data suggested that dysregulation of *FIP1L1* might could impair the stability or translation of mitochondrial-related mRNAs, indirectly affecting mitochondrial activity.

Taken together, our findings illustrate that integrating multi-omics data with systems genetics approaches enables deeper insights into how molecular mechanisms shape organism-level responses, such as mitochondrial stress regulation and lifespan determination, providing a foundation and resource for future aging research across different biological scales and models.

## Methods

### Bacterial and C. elegans strains

The Bristol strain (N2) and Hawaii were used as the wild-type strains were obtained from the *Caenorhabditis* Genetics Center (CGC; Minneapolis, MN). *E. coli* OP50 and HT115 strains were also obtained from the CGC. RNAi clones against *dmd-5*, *irx-1*, *nhr-25*, *fipp-1*, *F13A2.4*, *Y47D7A.12*, *Y47D7A.15*, *F32D1.7*, *neg-1* and *oac-12* were obtained from the Ahringer libraries or vidal library and verified by sequencing before use. Worms were cultured and maintained at 20°C and fed with *E. coli* OP50 on Nematode Growth Media (NGM) plates unless otherwise indicated.

### Lifespan measurements

Lifespan measurement was performed as previously described^62^. For both the RIAILs and wild-type N2 validation experiments, 5–10 L4-stage worms were transferred onto RNAi plates (containing 2 mM IPTG and 25 µg/mL carbenicillin) seeded with E. coli HT115, with or without 15 µg/mL doxycycline (Dox; Sigma, Cat. D9891). Once F1 progeny reached the L4 stage, worms were transferred to corresponding plates supplemented with 10 µM 5-fluoro-2’-deoxyuridine (5FU) to prevent progeny production. The L4 stage was defined as day 2 of life. Approximately 60 worms (for RAILs) or 80 worms (for validation) were used per condition and scored every other day. Worms were recorded as dead when unresponsive to gentle prodding with a platinum wire. Animals that went missing, displayed internal hatching, had vulval ruptures, or burrowed into the agar were censored. All validation experiments were independently repeated at least twice. Survival curves, log-rank p-values, and hazard ratios were analyzed using GraphPad Prism 9.

### Phenotyping by microfluidics

The phenotyping process was described in the previous study^33^. Briefly, the phenotypic readouts reflecting development, growth dynamics, fertility and reproduction of different RIAIL strains were collected with the SydLab robotic microfluidic-based platform developed by Nagi Bioscience, which allows high-content *C. elegans* screenings. A synchronized population of *C. elegans* was injected into microfluidic chips at the L1 larval stage. Worms were confined within dedicated microfluidic chambers and were continuously fed with freeze-dried *E. coli* OP50 solution. For the Dox condition, doxycycline was dissolved in bacterial solution at a concentration of 15 µg/mL, and worms were chronically exposed to Dox from the L1 stage until the end of the experiment (∼100 h post-worm injection into the microfluidic platform). The images of each chamber were recorded every hour for the whole duration of the experiment. At the end of the experiment, the collected images were processed by a set of software modules (also developed by Nagi Bioscience) based on machine learning algorithms, allowing a fully automated and standardized way for feature extraction and data analysis. The experiments were performed at 23°C.

The lmer function of the lmerTest (version 3.1-3) R package was used to adjust for batch effects in data collection. The following formula was used “value ∼ (1|batch/channel)” with default parameters.

### Sample collection for RNA-seq, proteomics, lipidomics, and metabolomics analyses

Worms of each RIAIL strain were cultured on plates seeded with *E. coli* OP50, then worm eggs were obtained by alkaline hypochlorite treatment of gravid adults. A synchronized L1 population was obtained by culturing the egg suspension in sterile M9 buffer overnight at room temperature. Approximately 2000 L1 worms of each RIAIL were transferred onto plates with or without 15 µg/mL Dox seeded with *E. coli* HT115. L4 worms were harvested after 2.5 days with M9 buffer and washed for three times. Worm pellets were immediately submerged in liquid nitrogen for snap-freezing and stored at −80°C until use.

### RNA extraction and RNA-seq data analysis

On the day of the RNA extraction, 1 mL of TriPure Isolation Reagent was added to each sample tube. The samples were then frozen and thawed quickly eight times with liquid nitrogen and a 37°C water bath to rupture worm cell membranes. A column-based kit from Macherey-Nagel was applied to extract total RNAs. RNA was sequenced by BGI with the BGISEQ-500 platform. The quality of the reads was then verified using FastQC (version 0.11.9)^58^. Low-quality reads were removed, and no trimming was needed. Alignment was performed against the worm genome (WBcel235 sm-toplevel) following the STAR (version 2.73a) manual guidelines^59^. Read counts were used for further analysis.

### Lipid extraction

All reagents were kept on ice, and samples were maintained at ≤ 4°C during the extraction procedure. A metal bead was added to each sample. Next, 500 µL M1 (MTBE:MEOH = 3:1, v:v) was added to each tube and vortexed for 2 min. 325 µL M2 (H2O:MEOH = 3:1, v:v) was added to each tube. Samples were vortexed briefly. Then, samples were flash-frozen in liquid nitrogen and thawed on ice. This step was done three times to facilitate cell breakage. Samples were transferred to a bead-beater and shaken at 1/25 s frequency for 5 min, and this process was done three times. The samples were then centrifuged for 10 min at 12,500 g at 4°C. For downstream lipid analysis, 200 µL of the organic layer (upper phase) was transferred to a glass autosampler vial and dried by vacuum centrifugation. For downstream metabolomic analysis, 200 µL of the aqueous layer (lower phase) was transferred to glass autosampler vials and dried by vacuum centrifugation. The remaining protein pellets on the bottom were kept on ice until further digestion.

Once dried, organic extracts intended for lipid analysis were resuspended in 100 µL 65:30:5 Isopropanol:Acetonitrile:Water and vortexed for 20 s prior to analysis by Liquid chromatography–mass spectrometry (LC-MS). Aqueous extracts intended for metabolomic analysis were resuspended in 50 µL 1:1 Acetonitrile:Water and also vortexed for 20 s prior to analysis by LC-MS.

### Protein digestion

Remaining protein pellets on the bottom were washed with 1 mL acetonitrile (ACN) and centrifuged at 10 kg for 3 min at 4°C. Supernatant ACN was aspirated and the protein pellets sit for 10-15 min at room temperature, or vacuum dried briefly to dry up the liquid in the bottom of the tube. 300 μL lysis buffer (8M urea with 100 mM tris(2-carboxyethyl)phosphine, 40 mM chloroacetamide and 100 mM tris (pH = 8.0) was added to each sample and vortexed till the protein pellets were fully dissolved. 5 μg LysC was added to each sample with protein:enzyme ratio 70:1 (digestion lasted overnight at room temperature). Trypsin at 70:1 protein:enzyme was added to each sample after diluting the lysis buffer to 2 M urea, and digestion lasted for six hours at room temperature. Desalting was carried out with 96-well desalting plates. A blank well between any two samples was reserved to avoid cross-contamination. Desalting started with equilibrating the desalting wells with 1 mL 100% ACN, followed by 1 mL 0.2% FA. Acidified peptide mixture was loaded to the 96-well desalting plate, followed by a 2 mL 0.2% FA wash. Peptides were eluted into a 96-well collection plate with 600 μL 80% ACN with 0.2% FA. Peptides were vacuum dried down and stored in −80°C freezer until resuspension with 0.2% FA. After resuspension, peptide concentration was measured using a quantitative colorimetric peptide assay.

### LC-MS setup

#### Proteomics

Peptides were separated on an in-house prepared high pressure reversed phase C18 column. Briefly, a 75-360 μm inner-outer diameter bare-fused silica capillary was packed with 1.7 μm diameter, 130 □ pore size, Bridged Ethylene Hybrid C18 particles (Waters) under high pressure of 25K psi to a final length of ∼40 cm. The column was installed onto a Thermo Ultimate 3000 nano LC and heated to 50°C for all runs. Mobile phase buffer A was composed of water with 0.2% FA. Mobile phase B was composed of 70% ACN with 0.2% FA. Samples were separated with a 120 min LC method: peptides were loaded onto the column for 13 min at 0.37 μL/min. Mobile phase B increased from 0 to 6% in 13 min, then to 53% B at 104 min, 100% B at 105 min, and held for 4 min at 100% B, decreased to 0% B at 110 min, and a 10 min re-equilibration at 0% B.

Eluting peptide fragments were ionized by electrospray ionization and analyzed on a Thermo Orbitrap Eclipse. Survey scans of precursors were taken from 300 to 1350 m/z at 240,000 resolution. Maximum injection time was 50 ms and automatic gain control (AGC) target was 1E6 ions. Tandem MS was performed using an isolation window of 0.5 Th with a dynamic exclusion time of 10 s. Selected precursors were fragmented using a normalized collision energy level of 25%. MS2 AGC target was set at 2E4 ions with a maximum injection time of 14 ms. Scan range was 150-1350 m/z. Scans were taken at the Turbo speed setting, and only peptides with a charge state of +2 or greater were selected for fragmentation.

#### Lipidomics

Extracted lipids were separated on an Acquity CSH C18 column (100 mm x 2.1 mm x 1.7 µm particle size; Waters) at 50°C using the following gradient: 2% mobile phase B from 0-2 min, increased to 30% B over next 1 min, increased to 50% B over next 1 min, increased to 85% over next 14 min, increased to 99% B over next 1 min, then held at 99% B for next 7 min (400 µL/min flow rate).

Column re-equilibration of 2% B for 1.75 min occurred between samples. For each analysis, 10 µL/sample was injected by autosampler. Mobile phase A consisted of 10 mM ammonium acetate in 70:30 (v/v) acetonitrile:milliQ H2O with 250 µL/L acetic acid. Mobile phase B consisted of 10 mM ammonium acetate in 90:10 (v/v) isopropanol:acetonitrile with 250 µL/L acetic acid.

The LC system (Vanquish Binary Pump, Thermo Scientific) was coupled to a Q Exactive Orbitrap mass spectrometer through a heated electrospray ionization (HESI II) source (Thermo Scientific). Source and capillary temperatures were 300°C, sheath gas flow rate was 25 units, aux gas flow rate was 15 units, sweep gas flow rate was 5 units, spray voltage was |3.5 kV| for both positive and negative modes, and S-lens RF was 90.0 units. The MS was operated in a polarity switching mode; with alternating positive and negative full scan MS and MS2 (Top 2). Full scan MS were acquired at 17,500 resolution with 1 x 106 AGC target, max ion accumulation time of 100 ms, and a scan range of 200-1600 m/z. MS2 scans were acquired at 17,500 resolution with 1 x 105 AGC target, max ion accumulation time of 50 ms, 1.0 m/z isolation window, stepped normalized collision energy (NCE) at 20, 30, 40, and a 10.0 s dynamic exclusion.

#### Metabolomics

Polar metabolites were separated on a Sequant ZIC®-pHILIC HPLC column (100 mm x 2.1 mm x 5 µm particle size) at 50°C using the following gradient: 95% mobile phase B from 0-2 min, decreased to 30% B over next 18 min, held at 30% B for 6 minutes, then increased to 95% B over next 1 min, then held at 95% B for next 8 min. The flow rate was 150 µL/min. For each analysis, 2 µL samples were injected and loaded onto the column. Mobile phase A consisted of 10 mM ammonium acetate in 10:90 (v/v) LC-MS grade acetonitrile: H2O with 0.1% ammonium hydroxide. Mobile phase B consisted of 10 mM ammonium acetate in 95:5 (v/v) LC-MS grade acetonitrile: H2O with 0.1% ammonium hydroxide.

The LC system (Vanquish Binary Pump, Thermo Scientific) was coupled to a Q Exactive HF Orbitrap mass spectrometer through a heated electrospray ionization (HESI II) source (Thermo Scientific). Source and capillary temperatures were 350°C, sheath gas flow rate was 45 units, aux gas flow rate was 15 units, sweep gas flow rate was 1 unit, spray voltage was 3.0 kV for both positive and negative modes, and S-lens RF was 50.0 units. The MS was operated in a polarity switching mode; with alternating positive and negative full scan MS and MS2 (Top 10). Full scan MS were acquired at 60K resolution with 1 x 106 AGC target, max ion accumulation time of 100 ms, and a scan range of 70-900 m/z. MS2 scans were acquired at 45K resolution with 1 x 105 AGC target, max ion accumulation time of 100 ms, 1.0 m/z isolation window, stepped NCE at 20, 30, 40, and a 30.0 s dynamic exclusion.

### Multi-Omics Data analysis

#### Proteomics

LC-MS files for proteomics were searched in Maxquant (version 2.0.3.1) against newly downloaded *C. elegans* proteome database from Uniprot. Parameters used in Maxquant can be found in the XML file named “mqpar”. Original outputs from Maxquant were inspected and potential contaminant proteins, protein groups that contain proteins identified with decoy peptide sequence, and those identified only with modification site were removed. LFQ intensities were used as the quantification metric.

#### Lipidomics

LC-MS files for lipidomics were processed using Compound Discoverer 3.1 (Thermo Scientific) and LipiDex. All peaks with a 1.4-23 min retention time and 100 Da to 5000 Da MS1 precursor mass were aggregated into compound groups using a 10 ppm mass tolerance and 0.4 retention time tolerance. Peaks were excluded if peak intensity was less than 2 x 106, peak width was greater than 0.75 min, signal-to-noise ratio was less than 1.5, or intensity was < 3-fold greater than blank. MS2 spectra were searched against an in-silico generated spectral library containing 35,000 unique molecular compositions of 48 distinct lipid classes^63^. Spectra matches with a dot product score > 500 and a reverse dot product score > 700 were retained for further analysis. Lipid MS/MS spectra that contained < 75% interference from co-eluting isobaric lipids, eluted within a 3.5 median absolute retention time deviation (M.A.D. RT) of each other, and were found within at least 4 processed files were used for identification at the individual fatty acid substituent levels of structural resolution. If individual fatty acid substituents were unresolved, then identifications were made with the sum of the fatty acid substituents. Peak intensities were normalized with the peptide amount to correct for different amounts of starting materials across the RIAIL panel.

#### Metabolomics

LC-MS files for metabolomics were processed using Compound Discoverer 3.3 (Thermo Scientific) in a discovery mode. All peaks between 0 and 22 min retention time and 0 Da to 5000 Da MS1 precursor mass were grouped into distinct chromatographic profiles (i.e., compound groups) and aligned against the reference file (the QC file running in the middle of all the files). Profiles not reaching a minimum peak intensity of 5 x 104, a maximum peak-width of 3 min, a signal-to-noise (S/N) ratio of 1.5, and a 5-fold intensity increase over blanks were excluded from further processing. Profiles having fewer than five points across the peak were also excluded. Element compositions were predicted with 5 ppm mass tolerance based on MS1 precursor mass. Precursors were matched to compounds by searching against databases including Biocyc, Human metabolome database and KEGG. MS/MS spectra were searched against mzcloud (Thermo Scientific), containing 19,503 unique molecular compositions, mzVault libraries including in-house curated MS2 spectra of 151 standards, 598 polar compounds from Bamba lab, Fiehn lab HILIC library of 3061 entries, KI-GIAR zic HILIC library of 814 entries, and six other libraries from MassBank of North America.

The resulting features were filtered based on the peak quality rating, and only the features that had peak rating greater than 4.0 (on the scale of 0-10) in more than one-third of the samples were kept for further analysis. Compound annotation was done manually by examining the composition, MS2 spectrum and retention time match. Peak intensities were also normalized with the peptide amount to correct for different amounts of starting materials across the RIAIL panel.

### Survival analysis and lifespan traits extraction

Lifespan data was analyzed using the survfit function of the R package survival (version 3.5-8) with formula: survival::Surv(Age_of_death, status) ∼ Strain + Dox exposure. Parental strains (N2 and CB4856) lifespan from each batch was compared to check for possible batch effects. No batch correction was performed. The quantile function was used to obtain the average lifespan as well as the 25%, 50% and 75% mortality.

### Differential expression analysis

Transcriptome: Normalised effective library sizes were calculated by TMM (trimmed mean of M-values). The voom function of the Limma R package (version 3.60.0)^64^ was applied to transform gene counts for linear modeling with precision weights.

#### Proteome

Log_2_ transformed proteomics data was then used in further analysis.

#### Metabolome and Lipidome

Normalized metabolomic and lipidomic datasets were calculated using variance stabilization normalization (Vsn) by vsn R package (version 3.72.0)^65^. The differential expression analysis for transcriptome, proteome, metabolome, and lipidome was performed using the R package Limma (version 3.60.0)^64^.

### Weighted gene co-expression network analysis (WGCNA)

WGCNA R package (version 1.72-5)^66^ was applied to construct a co-expressed gene network using the transcriptomic data under control and Dox conditions. Pearson correlation was performed to evaluate the correlations between all pairs of genes across all RIAILs. The pickSoftThreshold function was then applied to identify the best soft-thresholding power of 5 with networkType equal to signed hybrid, blockSize equal to 25000 and corFnc equal to bicor. The co-expression network is finally constructed with parameters (networkType = “signed hybrid”, minModuleSize = 30, reassignThreshold = 1e-6, mergeCutHeight = 0.15, maxBlockSize = 25000) using the calculated correlation coefficient of all pairs of genes. The eigengenes of each module were used for further analyses.

### Multi□omics factor analysis (MOFA)

Normalized proteomic, metabolomic, and lipidomic datasets were used to train MOFA model using the MOFA2 R package (version 1.14.0)^44^ with maxiter parameter 50,000 to ensure convergence and 10 factors and the remaining default parameters.

### Cox regression analysis

To estimate the association between lifespan-related traits and phenotypic traits, co-expressed modules, or multi-omics factors, the hazard ratio of a Cox regression model was estimated with R package survival^67^ using the following formula: survival::coxph(survival::Surv(lifespan-related traits) ∼ factor + treatment. *Geneset enrichment analysis (GSEA) and Over-representation analysis (ORA)*

Proteomic data: genesets used in GSEA and ORA were downloaded from wormbase (https://downloads.wormbase.org/releases/WS283/ONTOLOGY. The overlapped significantly altered genes/proteins upon Dox exposure were selected and then the ORA was performed using Clusterprofiler R package (version 4.2.2)^68^. Proteins were also ranked based on weights contributed to the factor 9 obtained from MOFA and GSEA was performed by Clusterprofiler R package (version 4.2.2)^68^.

Metabolomic data: The significantly changed metabolites (BH-adjusted P value < 0.05) amd the ORA were performed using MetaboAnalystR Package (version 4.0.0)^69^.

### Human dataset analysis

Summary statistics for UKBB phenotypic data were assessed through IEUGWAS^70,71^ and that for MVP database^72^ were accessed through dbGaP under accession phs002453.v1.p1 through AgingX project (ID: 10143).

### Mendelian randomization analysis

Summary statistics was used to estimate the causal effects of the tissue-specific expression of candidate genes on various outcomes. Significant eQTLs of candidate genes in the skeletal muscle were selected and their effect sizes were obtained from the GTEx Portal^48^ on 2023-03-28 (v8, https://www.gtexportal.org/home/datasets). GWAS summary statistics for *FIP1L1* genetic variants-related phenotypic traits were obtained from the IEU OpenGWAS project^70,71^, using the ieugwasr package (version 0.1.5). In most cases, multiple GTEx eQTLs were available in the outcome GWAS to be used as instrumental variables (IVs). Their effects were combined using a PCA-based approach^73^ to obtain independent IVs explaining 99% of the genetic variance, after which the causal effect was estimated through inverse-variance weighting MR using the TwoSampleMR R package^74^. In cases where only a single IV was available in both GTEx and the outcome GWAS (or if a single PC was retained), we used the Wald ratio method in TwoSampleMR^74^.

Because the effect sizes from GTEx^48^ do not have a direct interpretation or unit, the causal effect sizes are not representative, however the directionality of the effect remains informative. The scale of these QTL effects remains constant per gene, however, which enables their combination using the above-mentioned PCA-based approach^73^.

### Fluorescent image for assessing the UPR^mt^ activation

RNAi bacteria were cultured overnight in lysogeny broth (LB) medium containing 100 mg/mL ampicillin at 37°C. Then the bacteria were concentrated five times and seeded onto RNAi plates. Random L4/young adult *hsp-6p::gfp* worms were picked onto the RNAi bacteria-seeded plates and cultured at 20°C until their progenies reached the young adult stage. 8 worms were then randomly picked in a drop of 40 mM tetramisole (Cat. T1512, Sigma) and then aligned on an empty NGM plate. Fluorescent images, with the same exposure time for each condition, were captured using a Nikon SMZ1000 microscope.

### Real-time quantitative PCR (RT-qPCR)

For qRT-PCR, N2 worms were cultured and total RNA was extracted. cDNA synthesis was performed using the Qiagen Reverse Transcription Kit (205314) from the extracted RNA samples. The qPCR was then conducted with the Roche Light Cycler 480 SYBR Green I Master kit (Cat. 04887352001). The specific primers utilized are detailed in the key resources table, with *cdc-42* and *cdc-42* primers serving as housekeeping controls **(Table S1)**.

### Cell culture and drug treatment

HEK293T cells (Cat. CRL-3216) were obtained from ATCC. All cell lines were validated to be free of mycoplasma contamination and maintained in Dulbecco’s modified Eagle’s medium containing 4.5 g glucose per liter and 10% fetal bovine serum. Transfection was performed with the Lipofectamine™ RNAiMAX Transfection Reagent. SiRNA for human FIP1L1 is GAAACUGCCCUUCCAUCUA. The compounds used for treatment of cells were Doxycycline (Cat. D9891, Sigma) with 30□μg/mL.

### Western blot assay

Proteins were extracted with Radioimmunoprecipitation Assay (RIPA) buffer supplied with protease and phosphatase inhibitors, as described previously^15^. For western blotting, the antibodies used were: P-EIF2α (Cat. 3597, CST, 1:800), Tubulin (Cat. T5168, Sigma, 1:2000), ASNS (Santa Cruz, Cat. sc-365809, 1:1000), EIF2α (Cat. 9722, CST, 1:1000), ATF4 (Cat. 11815, CST, 1:1000), FIP1L1 (C-10) (Santa Cruz, Cat. sc-398392, 1:500).

## Data availability

The RNA-seq data have been deposited in the National Center for Biotechnology Information Gene Expression Omnibus database (accession no. GSE300044). The mass spectrometry raw files have been deposited with the MassIVE database (accession no. MSV000088622 contains founder strain proteomics data and MSV000089880 contains the multiomics data.

## Acknowledgments

We thank the Andersen lab and the *Caenorhabditis* Genetics Center for providing the *C. elegans* strains. We thank all members of the Johan Auwerx and Kristina Schoonjans laboratories for helpful discussions. The work in J.A.’s group was supported by grants from the EPFL, the European Research Council (ERC-AdG-787702), and the Swiss National Science Foundation (SNSF 31003A_179435, Sinergia CRSII5_202302, and SNSF-IZLCZ0-206069) and a GRL grant of the National Research Foundation of Korea (NRF 2017K1A1A2013124). The work in J.J.C.’s lab was supported by P41GM108538 (J.J.C.) and R35GM118110 (J.J.C.) from the National Institutes of Health. A.W.G. was supported by the United Mitochondrial Disease Foundation (PF-19-0232), Amsterdam UMC Postdoc Career Bridging Grant, Horizon-MSCA-PF-EF-2022 (101108082), and AGEM Talent Development Grant (2023). T.Y.L. was supported by the Human Frontier Science Program (LT000731/2018-L). Work in the Houtkooper group is financially supported by the Velux Stiftung (no. 1063), an NWO-Middelgroot grant (no. 91118006) from the Netherlands Organisation for Scientific Research (NWO), and a Longevity Impetus grant from Norn Group. W.L. is supported by the CSC (China Scholarship Council). H.L. was supported by research grants from Xi’an Jiaotong University and the National Natural Science Foundation of China (82300951).

## Author contributions

A.W.G., X.L., and J.A. conceived the project. A.W.G., E.K., Y.Z., T.Y.L. collected and measured all the measurements of RIAILs. W.L., Z.W., L.L. and L.P. performed the validation experiments. X.L. performed data analysis with great help from Y.Z., E.K., K.A.O., L.M., M.C., W.L., A.W.G., and H.L. for confirmation. J.A. A.W.G., R.H.H., and J.J.C. supervised the study. X.L., A.W.G., and J.A. wrote the manuscript with contributions from all the co-authors.

---

**Figure S1:**
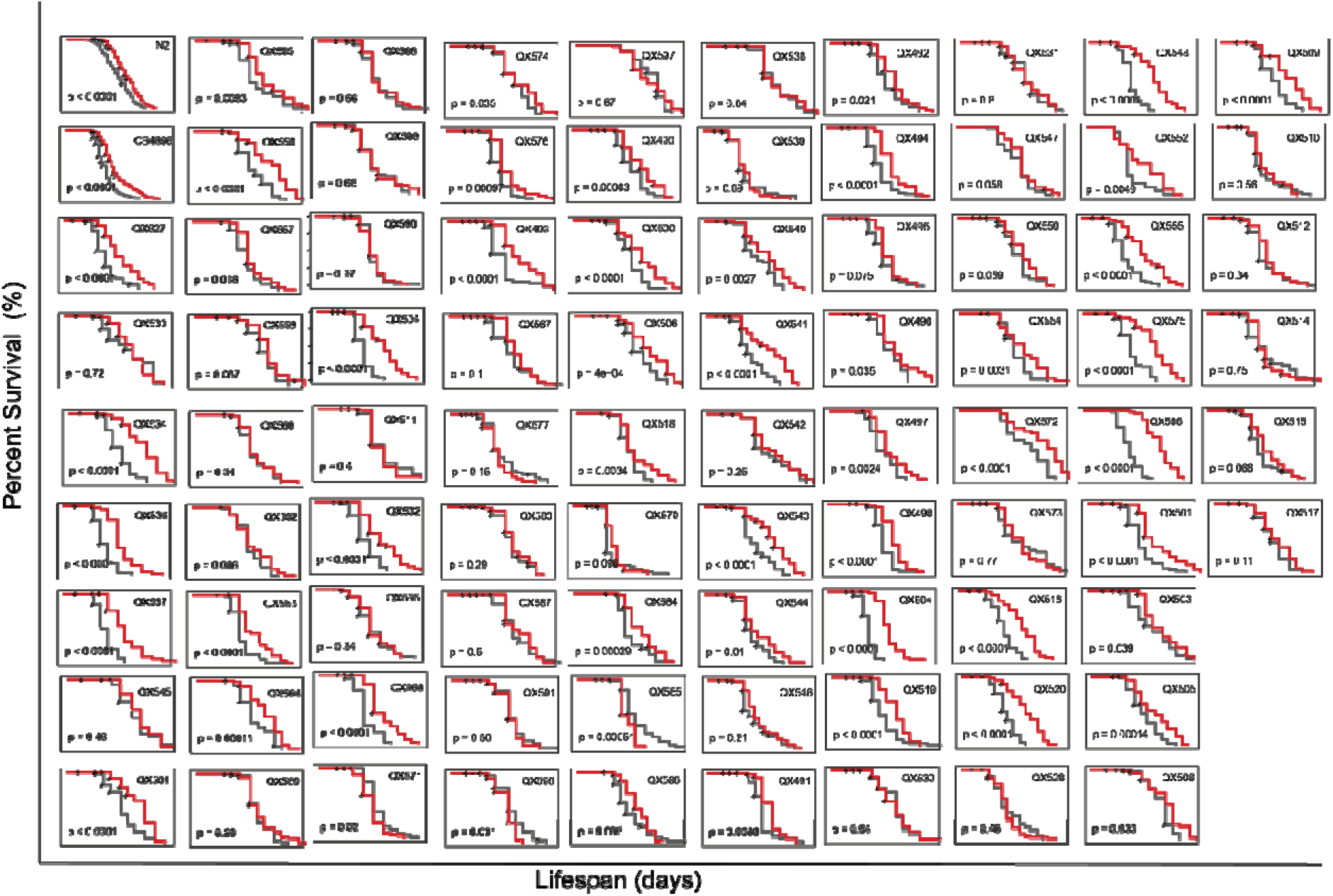
Lifespans of 85 RIAILs exhibit large variations at both control and Dox-treated conditions. Lifespans of 85 RIAIL strains. Control condition: black lines; Dox treated: red lines. Statistical analysis was performed by log-rank analysis (*p<0.05, **p<0.01, ***p<0.001, ****p<0.0001, ns. not significant).

**Figure S2:**
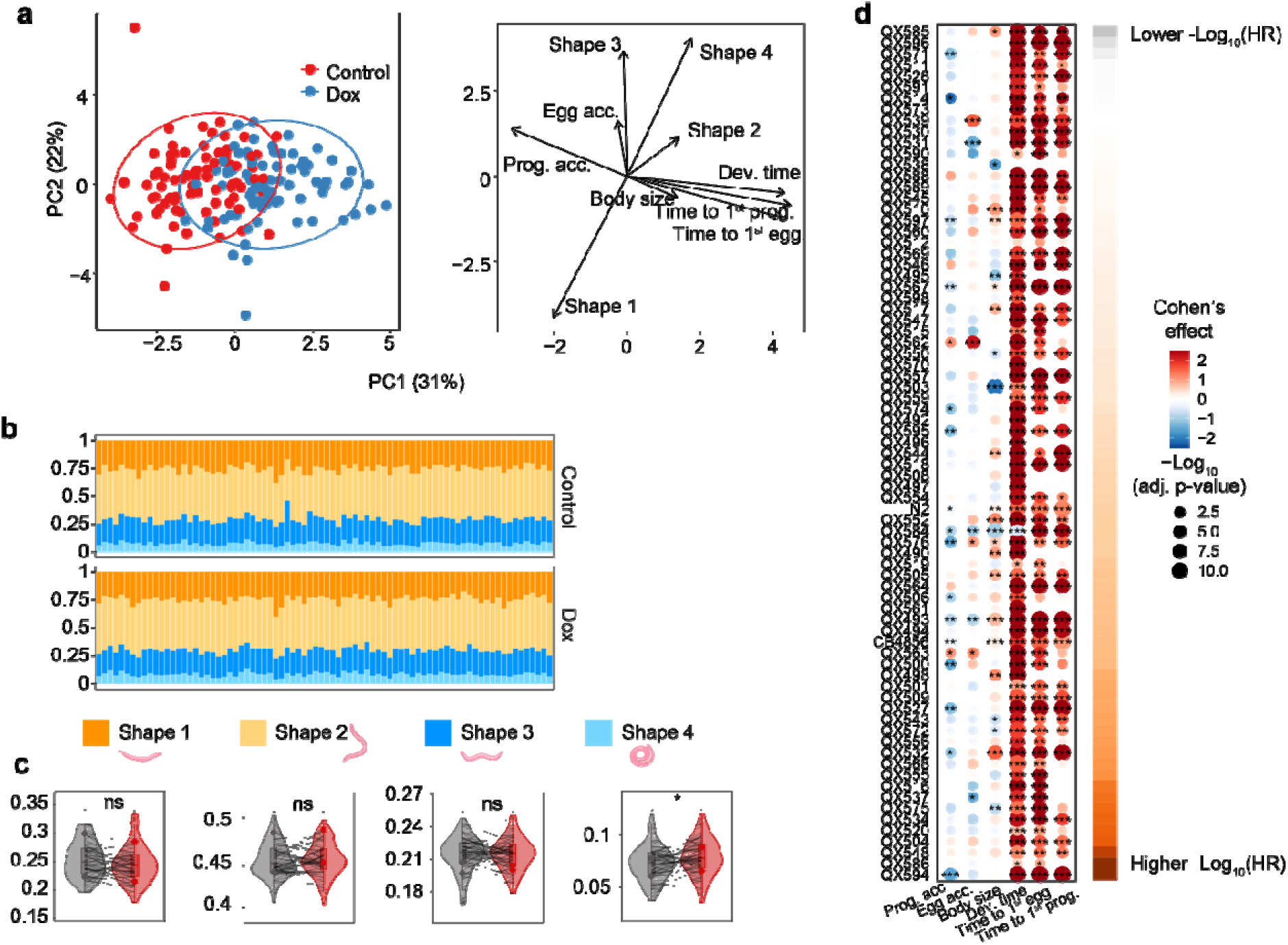
Dox effect on physiological phenotypes, transcriptome, and proteome. **(a)** PCA analysis showing the Dox effect on 10 clinical phenotypes and indicating the main affected phenotypes. **(b)** Percentage of time that a RIAIL strain spent in different shapes at control (upper) and Dox-treated (bottom) conditions. **(c)** Violin plots of the percentage of time that RIAIL strains spent in different shapes under control (black) and Dox (red) conditions. P-values represent the comparison of each trait between the control and Dox calculated using two-tailed Student’s t-test. Lines indicate the changes of each phenotype in each strain upon Dox. **(d)** Dot plot shows the changes of physiological traits upon Dox among RIAILs. Strains were ranked by lifespan extension induced by MSR (colored in orange). The Dox effect on these traits was evaluated by Cohen’s effect and represented by color. -Log_10_(adj.P.Val) i shown by dot size and indicated as * -Log_10_(adj.P.Val) < 0.05, ** -Log_10_(adj.P.Val) < 0.01, *** - Log_10_(adj.P.Val) < 0.001.

**Figure S3:**
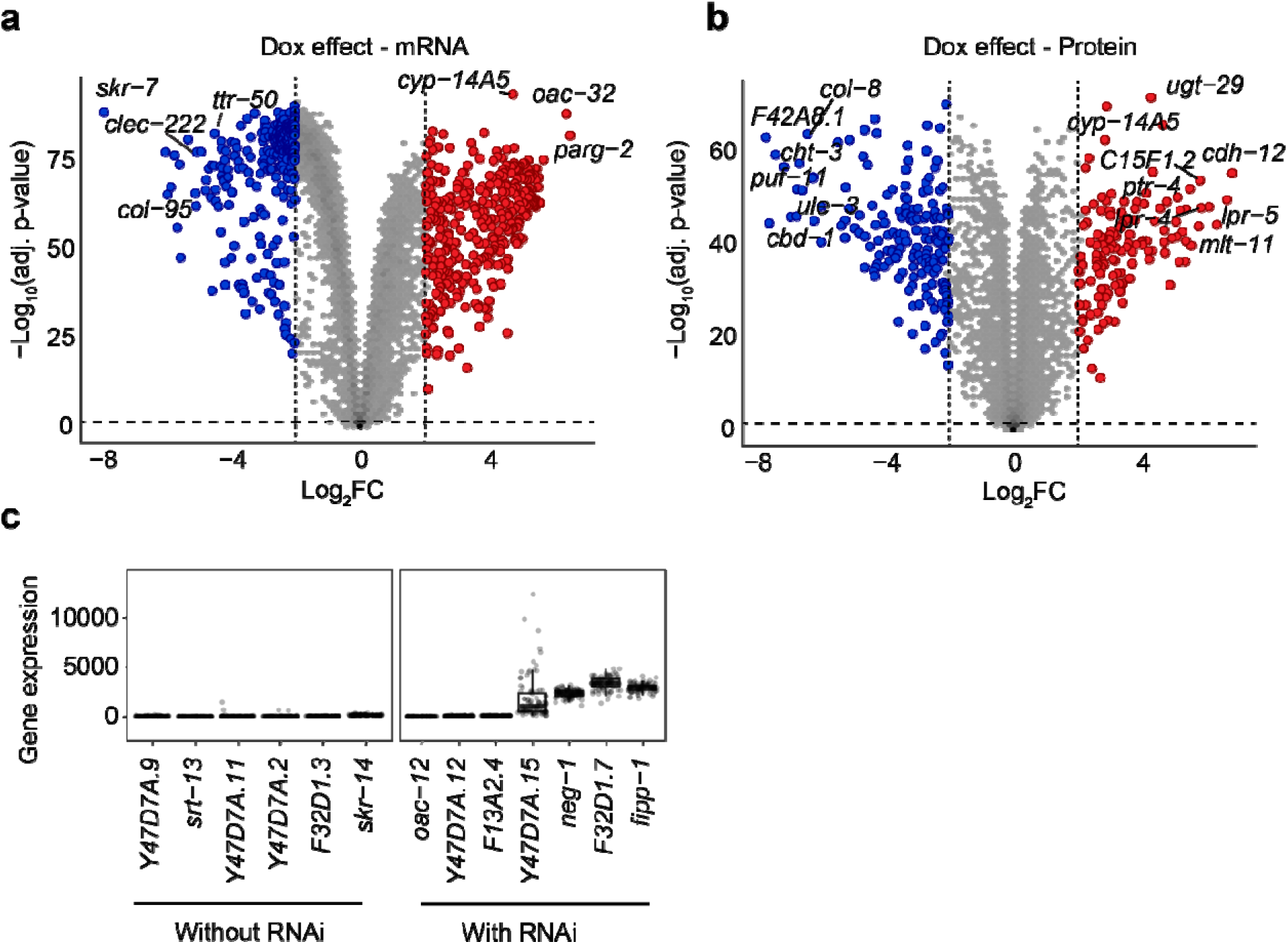
Dox effect on transcriptome, and proteome. **(a, b)** Volcano plots showing the Dox effect on transcriptome **(a)** and proteome **(b)**. Up- and down-regulated genes/proteins are indicated by red and blue, respectively. **(c)** Boxplot show that the candidate genes identified from the QTL peak with corrected RNAi have higher gene expression than those without RNAi or corrected RNAi.

**Figure S4:**
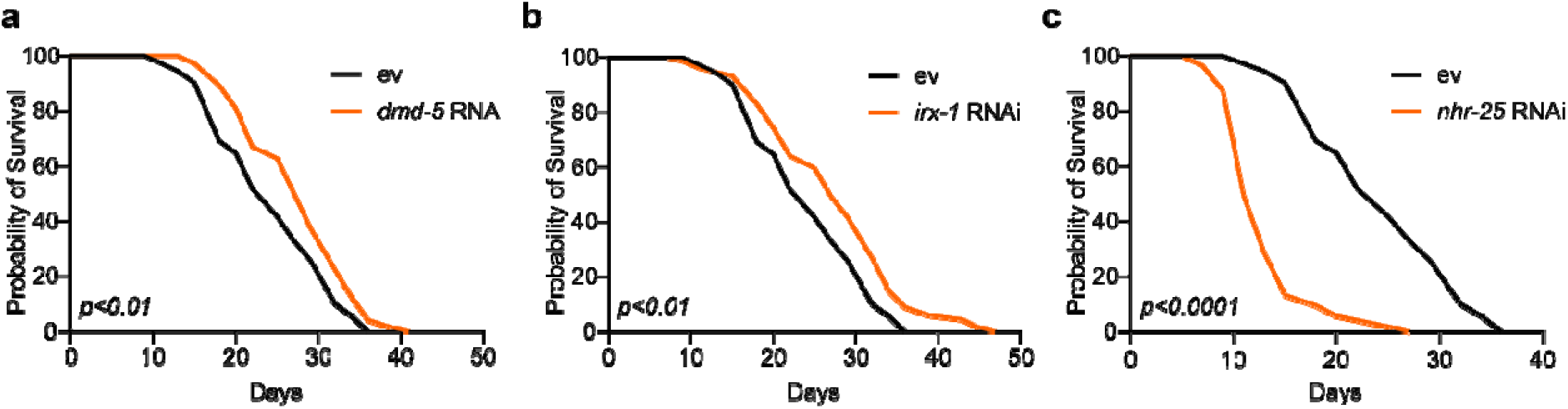
The effects of three identified longevity-related transcription factors. (**a-b**) RNAi of *dmd-5* (**a**) and *irx-1* (b) prolonged worm lifespan. (**c**) RNAi of *nhr-25* shortened the lifespan of *C. elegans.* Statistical analysis was performed by log-rank analysis.

**Figure S5:**
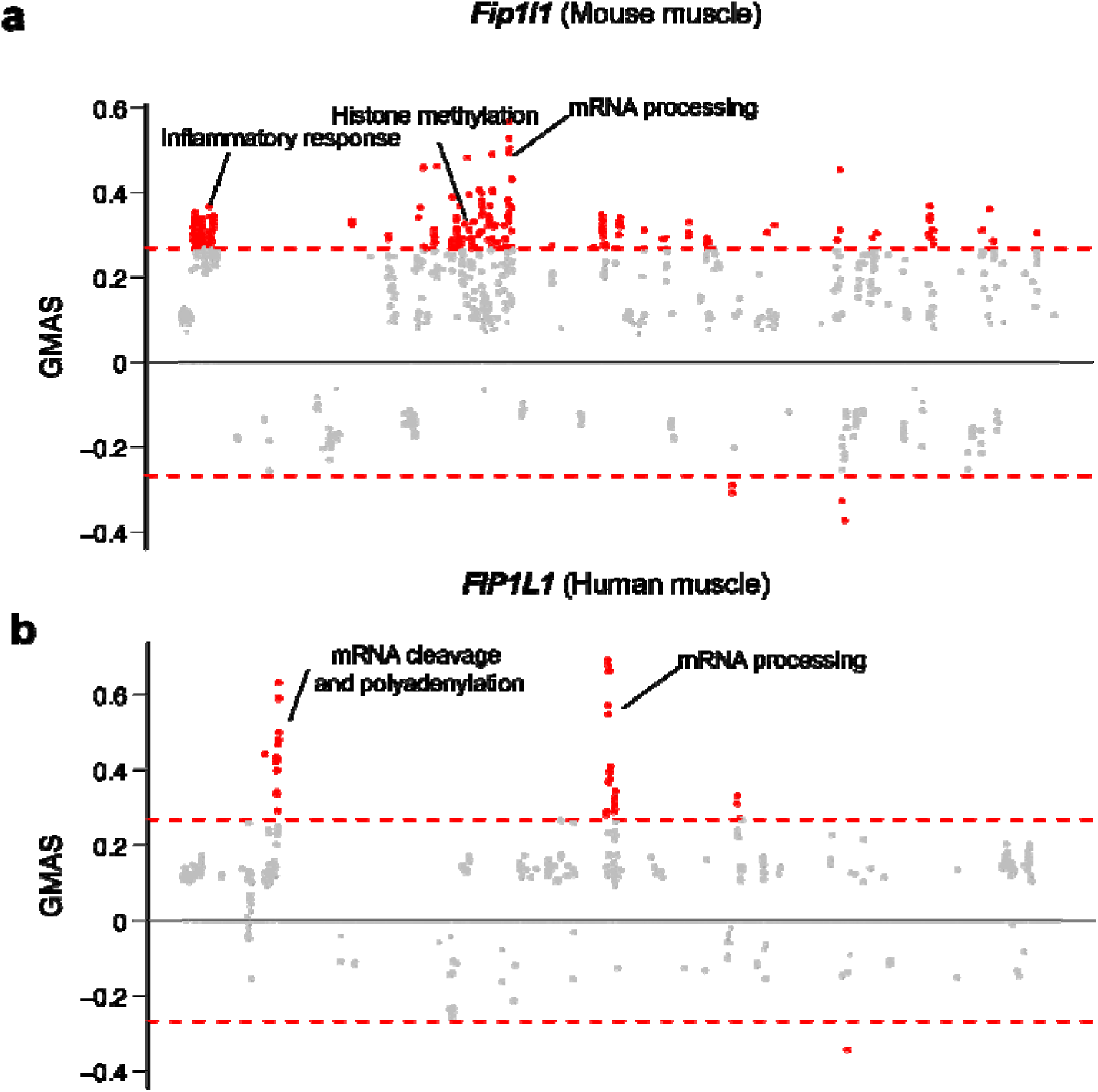
Exploring the biological function of FIP1L1 in mice and humans. (**a-b**) Manhattan plot showing the associated gene expression modules of *Fip1l1* in mouse **(a)** and human muscles **(b)**. Data were retrieved from https://systems-genetics.org/gmad^75^. The threshold is represented by the red dashed line (absolute gene-module association score ≥ 0.268). Terms above the threshold are identified as the significant associated terms. GO terms or gene modules are ranked by similarity.

**Table S1:**
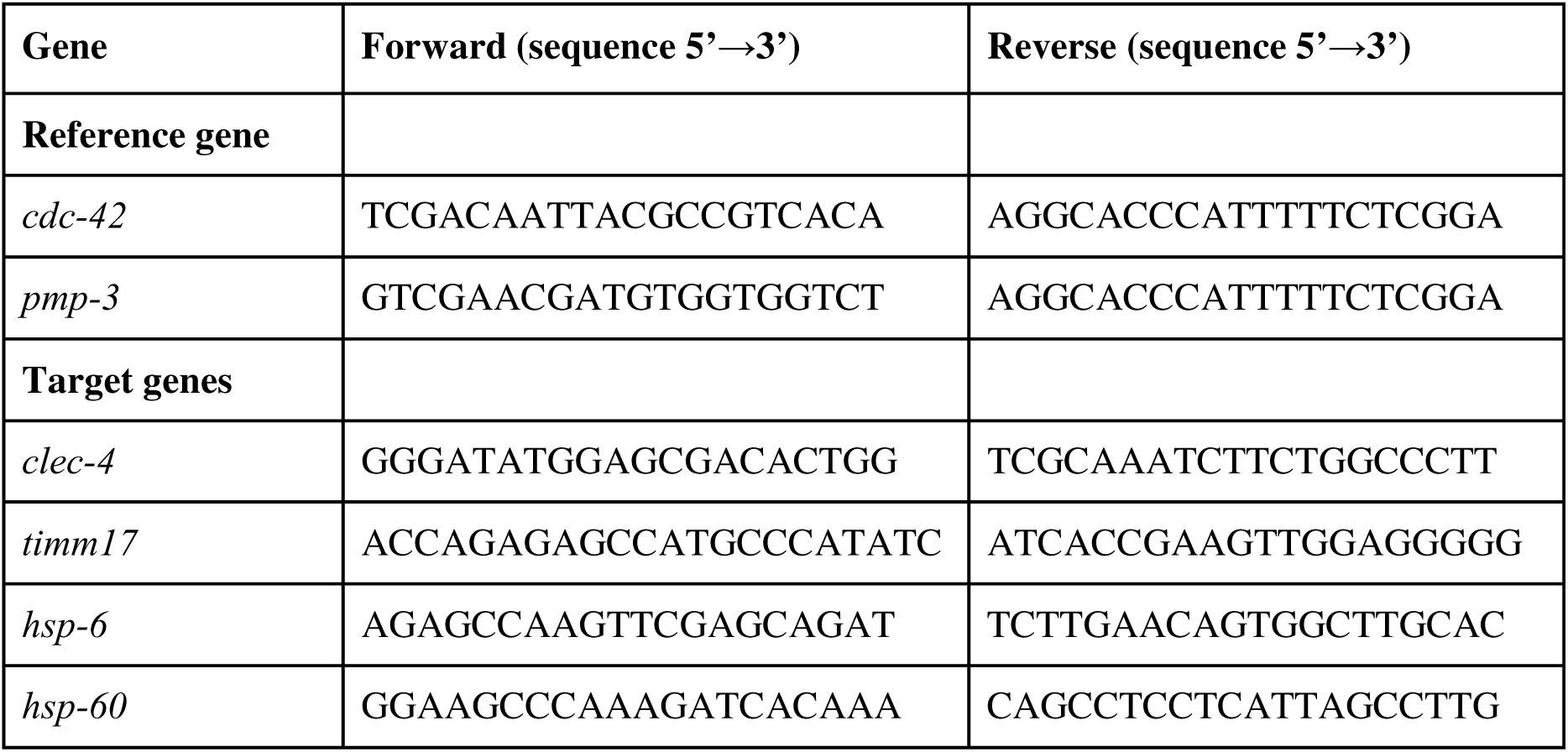
Primers used for *C. elegans* qPCR.

